# Neuronal identity is maintained in the adult brain through KAT3-dependent enhancer acetylation

**DOI:** 10.1101/836981

**Authors:** Michal Lipinski, Rafael Muñoz-Viana, Beatriz del Blanco, Juan Medrano-Relinque, Angel Marquez-Galera, Jose M. Carames, Andrzej A. Szczepankiewicz, Jordi Fernandez-Albert, Carmen M. Navarrón, Roman Olivares, Grzegorz M. Wilczynski, Santiago Canals, Jose P. Lopez-Atalaya, Angel Barco

## Abstract

Very little is known about the mechanisms responsible for maintaining cell identity in mature tissues. The paralogous type 3 lysine acetyltransferases (KAT3) CBP and p300 are both essential during development, but their specific functions in nondividing differentiated cells remains unclear. Here, we show that when both proteins are simultaneously knocked-out in excitatory neurons of the adult brain, the mice express a rapidly progressing neurological phenotype associated with reduced dendritic complexity and electrical activity, the transcriptional shutdown of neuronal genes, and a dramatic loss of H3K27 acetylation and pro-neural transcription factor binding at neuronal enhancers. The neurons lacking both KAT3 rapidly acquire a molecularly undefined fate with no sign of dedifferentiation, transdifferentiation or death. Restoring CBP expression or lysine acetylation reestablished neuronal-specific transcription. Our experiments demonstrate that KAT3 proteins act as *fate-keepers* in excitatory neurons and other cell types by jointly safeguarding chromatin acetylation levels at cell type-specific enhancers throughout life.

## Introduction

Neuronal identity is established through the action of a particular class of transcription factors (TFs), referred to as terminal selectors, that bind to cis-regulatory elements of terminal identity genes and regulate the establishment of neuron-specific gene programs ^1, 2^. Many of these terminal selectors are also required at later stages of development ^3^, which led to the idea that the identity of postmitotic neurons needs to be actively maintained throughout life by the action of the same TFs that controlled the last steps of differentiation ^4, 5^. However, the mechanisms responsible for such maintenance remain elusive. Chromatin-modifying enzymes are also involved in identity acquisition ^6, 7^, and although little is still known about their specific contribution, they are also likely involved in cell fate maintenance ^8^.

The KAT3 family of transcriptional co-activators comprises the CREB binding protein CBP (*aka* KAT3a) and the E1A binding protein p300 (*aka* KAT3b) ^9^. Both proteins have a lysine acetyltransferase (KAT) catalytic domain and interact with numerous other proteins, including histones, TFs, the RNA polymerase II complex (RNAPII), protein kinases and other chromatin-modifying enzymes ^10^ that also are substrates of the CBP/p300 catalytic activity ^11^. Consistent with this central function, KAT3 proteins play a critical and dose-dependent role during neurodevelopment ^12^. Deletions and inactivating mutations in KAT3 genes cause early embryonic death and neuronal tube closure defects ^13^, whereas hemizygous mutations cause a syndromic disorder associated with intellectual disability known as Rubinstein-Taybi syndrome (RSTS) ^14, 15, 16^. After birth, CBP loss in postmitotic forebrain neurons impairs memory in specific tasks, but does not interfere with cell viability and animal survival ^17, 18, 19^. The consequences of p300 loss in adult neurons have been less explored, but the only study conducted so far revealed very mild defects ^20^. The relatively modest consequences of the postembryonic loss of either one of these central transcriptional regulators could indicate that their role is limited to development. Alternatively, since both proteins are still expressed in adult neurons, they could be functionally redundant and thus mutually compensate for each other loss.

To define KAT3 specific functions in the adult brain, we generated mice with inducible and restricted ablation of CBP and/or p300 in forebrain principal neurons. Comprehensive characterization of these strains revealed that while both proteins are individually dispensable for the normal function of mature neurons, their combined ablation in the adult brain had devastating and rapid consequences: dendrites retract, synapses are lost, and electrical activity and neuron-specific gene programs established during development are shutdown. This phenotype leads to severe neurological defects and premature death. Epigenomic screens and rescue experiments demonstrate that both KAT3 proteins are jointly essential for maintaining the identity of excitatory forebrain neurons (and likely other cell types) throughout life by preserving acetylation levels at cell type-specific genes and enhancers.

## Results

### Combined, but not individual, loss of CBP and p300 in adult excitatory neurons causes severe neurological deficits

To elucidate the neuronal roles of CBP and p300 in the adult brain, we selectively eliminated CBP, p300, or both KAT3 proteins in forebrain excitatory neurons of adult mice using the inducible Cre-recombinase driver CaMKIIα-CreERT2 (**Fig. 1a**). The three inducible forebrain-specific knockout strains (referred to as CBP-ifKO, p300-ifKO and dKAT3-ifKO, respectively) do not show any neurological symptom before gene(s) ablation (**Table S1**). After tamoxifen (TMX) treatment of 3-month old mice, loss of immunoreactivity was observed in virtually 100% of the pyramidal neurons in the CA1 and cortex, and granule neurons in the dentate gyrus (**Fig. 1b** and **S1a**), while brain regions in which the CaMKIIα promoter is not active, such as the cerebellum and the basal ganglia, were spared. Importantly, the loss of either one of these paralogous proteins did not affect the expression level of the other (**Fig. S1b**).

**Figure 1.**
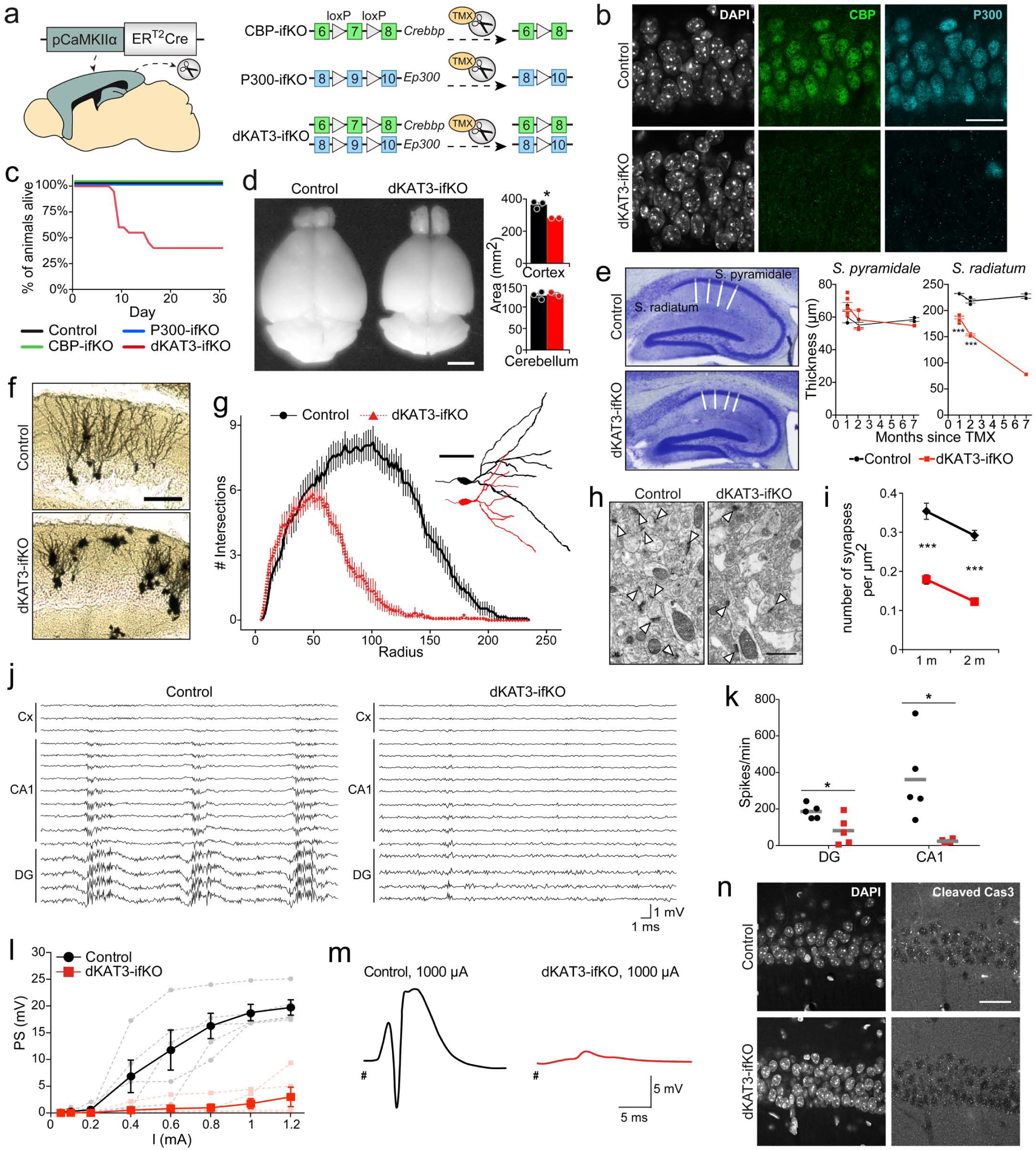
Loss of both KAT3 proteins causes severe neurological alterations. **a.** Genetic strategy for the production of inducible, forebrain-specific CBP, p300 and double KAT3 knockouts. **b.** Double immunostaining against CBP and p300 in the CA1 region. Scale: 100 µm. **c.** Survival of the three ifKO lines after TMX administration. (Control, n = 14; dKAT3-ifKO, n = 20). **d**. Left: Representative images of control and dKAT3-ifKO brains 2 months after TMX. Scale: 0.5 cm. Right: Quantification of cortical and cerebellar sizes (control, black bars, n = 3; dKAT3-ifKO, red bars, n = 2). Statistic: t-test. **e**. Left: Nissl staining of hippocampi 2 months after TMX. Right: Quantification of the thickness of the strata pyramidale and radiatum of control (1 m, n = 3; 2 m, n = 3; 7 m, n = 2) and dKAT3-ifKO mice (1 m, n = 6; 2 m, n = 2; 7 m, n = 1) at different time points after TMX. Points show separate observations. Statistic: two-way ANOVA. **f**. Representative images of Golgi staining in the dentate gyrus 1 month after TMX. Scale: 100 µm. **g**. Sholl analysis of DG neurons 1 month after TMX (32 neurons from 4 control mice; 38 neurons from 4 dKAT3-ifKOs). The right inset shows representative Neurolucida-reconstructed neurons. Scale: 50 µm. **h**. Electron microscopy images showing the stratum radiatum 1 month after TMX. Arrowheads indicate positions of the synapses. Scale: 1 µm. **i**. Number of synapses per µm^2^ in the stratum radiatum (average of 30 areas from 3 mice per condition). Statistic: t-test. **j**. Representative *in vivo* electrophysiological recordings of spontaneous activity across cortical and hippocampal layers. **k**. Firing frequency (spikes per minute) recorded in CA1 and DG. SUA: single unit analysis (n = 5). **l**. Population spike (PS) amplitude in DG after the application of increasing intensities in the perforant pathway (n = 5). Brighter-colored elements show the results for each animal separately. Statistic: t-test. **m.** Representative evoked potentials waveforms (PS) to 1 mA stimulation. **n**. Immunohistochemistry for cleaved Cas3 shows no sign of apoptosis in the CA1 subfield 1 month after TMX. Scale: 30 µm.

Consistent with the characterization of other forebrain-specific KO strains for CBP or p300 ^17, 18, 20^, CBP- and p300-ifKO mice had a normal lifespan and showed no overt neurological abnormalities (**Fig. 1c**). However, the situation was markedly different in dKAT3-ifKOs. When CBP and p300 were simultaneously removed in the forebrain of adult mice, the animals displayed a dramatic and rapidly progressing deteriorating phenotype (**Table S1** and **Fig. S1c-d**). In the first days following TMX administration, the mice were hyperactive and frequently froze in bizarre positions. Shortly afterwards, the same mice showed severe ataxia, and loss of the righting reflex, escaping response and tail-suspension-evoked stretching (**Movie S1**). Although different animals reached the terminal phenotype at different time points, all mice died within the first 2-3 weeks after TMX administration. Upon facilitating access to food and water, survival increased by about 40% within the first month after TMX administration (**Fig. 1c**), which suggests problems in self-alimentation. Remarkably, mice carrying just a single functional allele encoding either one of the two KAT3s did not die prematurely and showed normal reflexes (**Table S1** and **Fig. S1d-e**).

### Neurons lacking both KAT3 proteins display massive synaptic loss and reduced electrical activity, but do not die

Consistent with the severe neurological phenotype, the few dKAT3-ifKOs that survived for more than 2 months after TMX showed an obvious reduction of cerebral cortex volume, whereas other brain regions not targeted by gene deletion, such as the cerebellum, remained unaffected (**Fig. 1d**). Interestingly, histological analyses indicated that this reduction was fundamentally caused by a loss of neuropils rather than a loss of neuronal cells. For instance, in the hippocampus of dKAT3-ifKOs, the *stratum pyramidale* – a layer in the CA1 subfield occupied by the somas of pyramidal neurons – had a normal thickness and did not show any obvious sign of neurodegeneration (**Fig. 1e** and **Fig. S2a**) or gliosis (**Fig. S2b**) even several months after TMX treatment. In contrast, the *stratum radiatum* – a layer occupied by the basal dendrites of the CA1 neurons – was markedly thinner in dKAT3-ifKOs as soon as 1 month after TMX (**Fig. 1e**). In agreement with these quantifications, Golgi staining revealed a retraction of dendrites in the dentate gyrus (**Fig. 1f-g**) and electron microscopy (EM) analyses confirmed the massive loss of synapses in dKAT3-ifKOs (**Fig. 1h-i**). Furthermore, as early as two weeks after gene ablation, dKAT3-ifKOs implanted with multichannel electrodes displayed a dramatic reduction in both spontaneous (**Fig. 1j-k** and **S2c-d**) and evoked (**Fig. 1l-m** and **S2e**) electrical activity. Overall, these data demonstrate that the simultaneous loss of CBP and p300 alters neuronal morphology and impairs electrical responses leading to dramatic neurological defects.

Strikingly, these severe changes occur in the absence of cell death. TUNEL staining (**Fig. S3a**) and immunostaining against active caspase 3 (**Fig. 1n**) and the H2A.X variant (**Fig. S3b**), which respectively label cells undergoing apoptosis and suffering DNA damage, were all negative. EM images also show a largely normal *stratum pyramidale* in which neuronal nuclei did not present apoptotic bodies, although the nucleoplasm appeared clearer in dKAT3-ifKOs than in control littermates (**Fig. S3c-d**). To monitor the evolution of double knockout neurons beyond the limit imposed by the survival of dKAT3-ifKO’s, we infected the CA1 of the *Crebbp^f/f^::Ep300^f/f^*(dKAT3-floxed) mice with adeno-associated virus (AAV) expressing Cre recombinase under the human synapsin promoter (**Fig. S3e**). Immunostainings confirmed the efficient and complete elimination of CBP and p300 in granular neurons (**Fig. S3f**) in the absence of detectable cell death even two months after gene ablation (**Fig. S3g**).

### Maintenance of neuronal identity requires at least one KAT3 gene

To determine the molecular basis of the abovementioned phenotypes, we conducted a RNA-seq screen in the hippocampus of dKAT3-ifKOs and control littermates. Differential gene expression profiling revealed 1,952 differentially expressed genes (DEGs) in dKAT3-ifKOs, with a clear preponderance both in number and magnitude of gene downregulations (**Fig. 2a-b**, **S4a** and **Table S2**). Gene Ontology (GO) enrichment analysis indicated that these downregulations affect a large number of neuronal functions (**Fig. 2c**, blue bars). Hundreds of genes with neuronal functions such as genes encoding channels and proteins important for synaptic transmission were downregulated in the dKAT3-ifKO hippocampus, which explains the reduced neuronal firing and lack of electrical responses. Gene upregulation was much more restricted, including a modest inflammatory signature (**Fig. 2c**, red bars) but no activation of cell death pathways (**Fig. S4b**). In fact, several positive regulators of neuronal death were strongly downregulated in dKAT3-ifKOs (e.g., *Hrk;* **Fig. S4c**). Consistent with the survival of these cells, housekeeping genes remained largely unchanged (**Fig. 2b** and **S4d**). Immunodetection experiments for neuronal proteins like CaMKIV, NeuN and hippocalcin confirmed the dramatic loss of expression of neuronal proteins (**Fig. 2e** and **S4e**). Notably, the loss of neuronal markers expression was not detected in mice bearing a single functional KAT3 allele (**Fig. S4f**), indicating that this minimal gene dose is sufficient to preserve the active status of neuronal loci.

**Figure 2.**
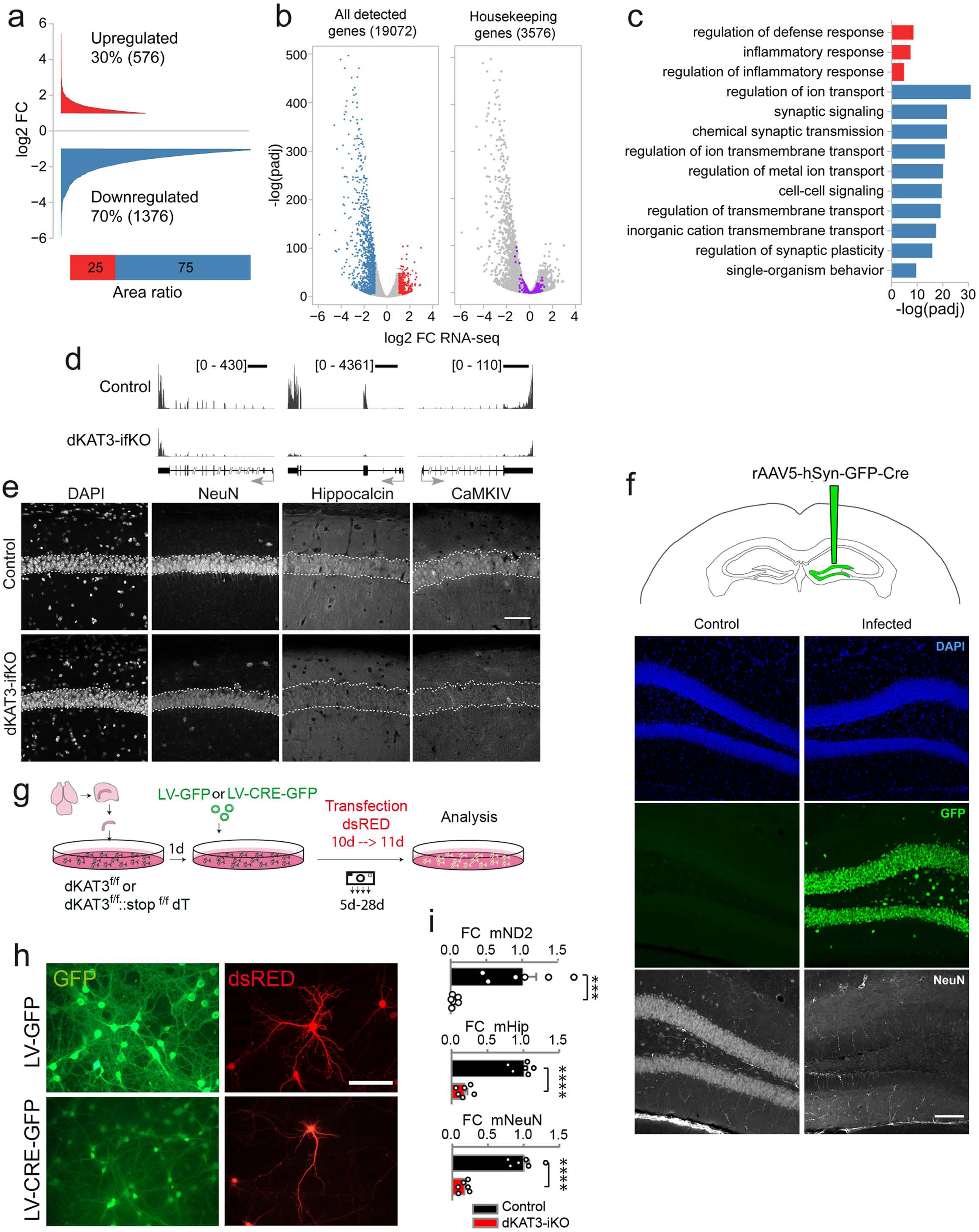
Hippocampal cells lacking KAT3 fail to express neuronal-specific genes. **a**. Cumulative graph showing the log2 fold-change value of DEGs in dKAT3-ifKOs (mRNA-seq, 1 month after TMX, n=3 per genotype). Upregulated genes are presented in red and downregulated genes in blue (p.adj < 0.05 and |log2FC| ≥ 1). The bottom bar graph compares the area in each set. **b**. Volcano plots of RNA-seq analysis. From left to right, we present all genes (left) and the subset of housekeeping genes listed in ^72^ (right). Grey: genes that are not significantly deregulated; red: upregulated genes; blue: downregulated genes; purple: housekeeping genes. **c**. The ten most enriched categories identified by Gene Ontology (GO) analysis on up-(red) and downregulated (blue) genes with a p.adj < 0.05 and |log2FC|≥1. The upregulated gene set only retrieved three categories. **d**. RNA-seq profiles for three representative neuronal-specific genes: *Rbfox3* (NeuN), *Hpca* and *Camk4*. Scale: 2 kb. **e**. Immunohistochemistry against the neuronal protein encoded by *Camk4*, *Rbfox3* (NeuN) and *Hpca*, Dashed line labels the position of the *stratum pyramidale* based on DAPI images. Scale: 60 µm. **f.** Cell autonomous loss of neuronal identity after double KAT3 ablation using a cre recombinase-expressing AAV. The virus was injected unilaterally into the DG of adult *Crebbp^f/f^::Ep300^f/f^* mice. Immunohistochemistry analysis shows the downregulation of NeuN only in the recombined (GFP^+^) side. **g.** Scheme representing the strategy to eliminate both KAT3 proteins in hippocampal PNCs from E17 dKAT3^f/f^ embryos. **h.** Representative images showing morphological changes in *Crebbp^f/f^::Ep300^f/f^*hippocampal neurons infected with LV-CRE compared with LV-GFP control. **i.** RT-PCR demonstrates decreased levels of *Neurod2* (ND2), *Hpca* (hippocalcin, Hip) and *Rbfox3* (NeuN) transcripts in dKAT3-KO PNCs.

To determine if these deficits were cell-autonomous, we analyzed the hippocampus of dKAT3-floxed mice monolaterally infected with Cre-recombinase-expressing AAVs. We observed a dramatic loss of neuronal marker expression only in the transduced hippocampus (**Fig. 2f**). Moreover, the cells depleted of KAT3 proteins completely failed to respond to kainic acid, a strong agonist of glutamate receptors, further confirming the loss of excitatory neuron properties (**Fig. S5**). Similarly, experiments in primary neuronal cultures (PNCs) produced from the hippocampi of *Crebbp^f/f^::Ep300^f/f^* embryos infected with a Cre-recombinase-expressing lentivirus (**Fig. 2g**) demonstrate that the cultured neurons do not die after simultaneous elimination of CBP and p300 (**Fig. S6a-b**). They do, however, show abnormal morphology (**Fig. 2h**), downregulated neuron-specific transcripts (**Fig. 2i**) and proteins (**Fig. S6c**) and severe hypoacetylation of KAT3 targets (**Fig. S6d**).

### Cells lacking KAT3 proteins acquire a novel, molecularly undefined fate

The altered morphology, electrophysiological properties and gene expression all suggest that the excitatory neurons rapidly lose their identity after the elimination of both KAT3 genes. To tackle this hypothesis, we compared the set of DEGs in dKAT3-ifKOs with transcriptome information for the different cell types in the adult mouse brain using single-cell RNA-seq (scRNA-seq) data from the mouse cortex ^21^ (**Fig. 3a**). Our analysis revealed that the genes typically expressed in CA1 and S1 pyramidal neurons were significantly downregulated in the hippocampus of dKAT3-ifKOs, whereas other cell-type specific transcriptional programs were unaffected except for a modest increase of microglia genes related to inflammation. Importantly, although identity loss is often associated with dedifferentiation (i.e., the regression to an earlier stage of differentiation), we did not detect an upregulation of stemness genes ^22, 23^, nor neuronal stem cell (NSC)- or neuroprogenitor (NPC)-specific gene expression ^24^ (**Fig. 3a and S7a-b**). Trans-differentiation can be also discarded because we did not detect the upregulation of the transcriptional signatures of other brain cell-types.

**Figure 3.**
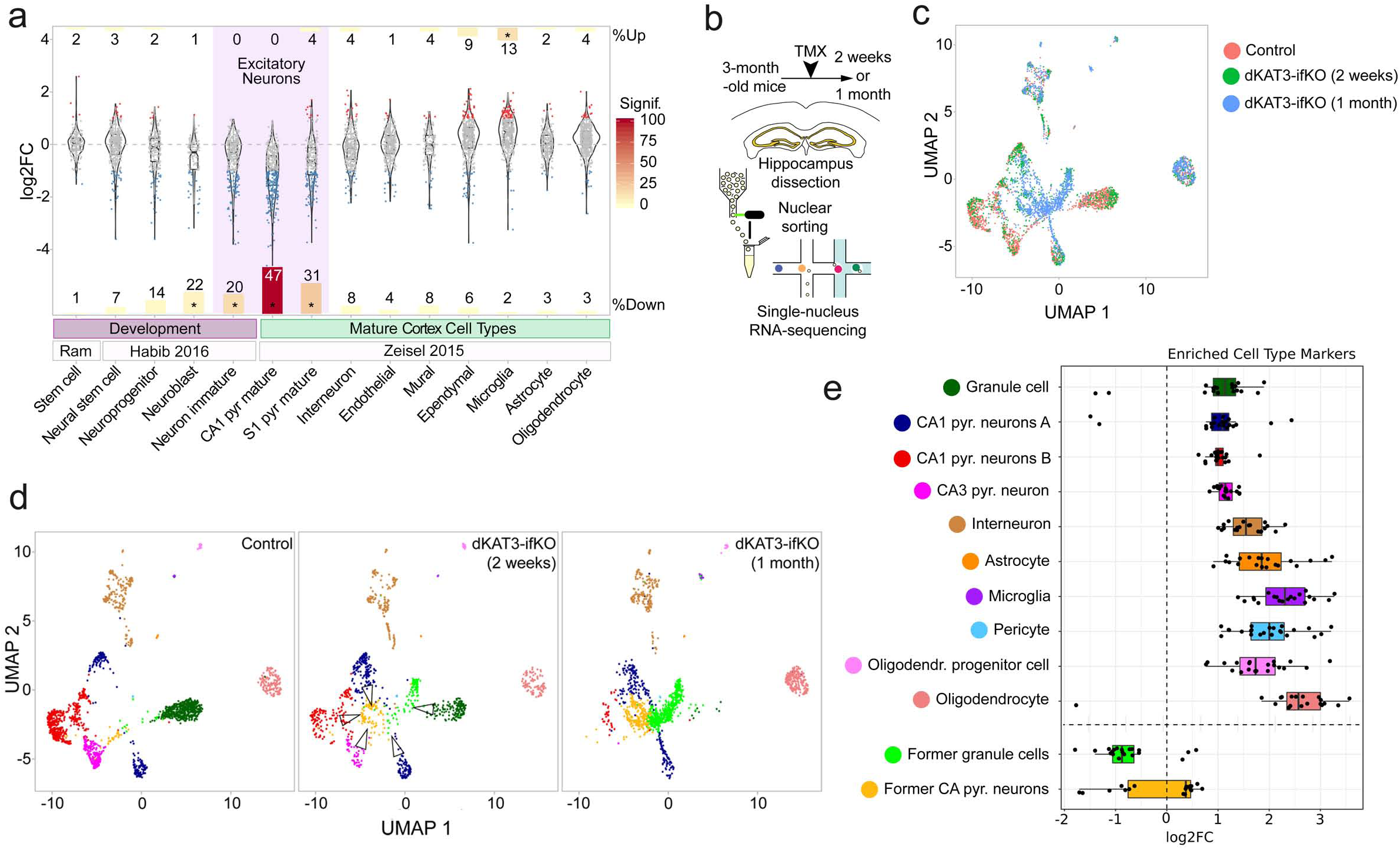
Cells lacking KAT3 proteins do not die or dedifferentiate, but acquire a novel, molecularly undefined fate. **a**. Violin plots show the change in expression of gene sets associated with (i) stemness ^23^; (ii) different neuronal differentiation stages (NSC: neural stem sell, NPC: neural progenitor cell, NB: neuroblast, IN: Immature neuron ^24^); and (iii) different cell types in the adult mouse cortex ^21^. Each dot represents a single gene. Bars are proportional to the percentage of up- or downregulated genes in each gene set. The significance of the enrichment resulting from a hypergeometric test (log10 of p.adj value) is indicated by the columns color. *: p < 0.05. **b.** Scheme of the single-nucleus RNA-seq experiment. **c.** UMAP plot of integrated analysis of snRNA-seq datasets from the dorsal hippocampus of dKAT3-fKOs and control littermates. **d.** UMAP plots showing identified populations in the hippocampus of control littermates (left) and dKAT3-ifKO mice 2 (center) and 4 (right) weeks after TMX treatment. Nuclei are colored by their classification label as shown in panel j. **e**. Box and whisker plot showing the expression of the top 20 markers for the ten cell types detected in the hippocampus of control mice (**Table S3**). The two bottom rows show the marker retrieved for the new cluster detected in dKAT3-ifKO mice 1 month after TMX. Intriguingly, these cells did not only stop to express granule and pyramidal neuron markers, they do not express either any marker that positively differentiate these cells from the other cell types (**Table S3**). Abbreviations: Pyr, pyramidal; Oligodendr, oligodendrocyte.

To determine more precisely the fate of excitatory neurons after losing their identity, we conducted single-nucleus RNA-seq analyses 2 and 5 weeks after TMX treatment (**Fig. 3b-c**, **S7c** and **Table S3**). We observed a progressive confluence of CA1/CA3 pyramidal neurons and dentate gyrus granule neurons in a common cell cluster depleted of neuronal type-specific markers (**Fig. S7d-g**) and in which no other distinctive marker appears (**Fig. 3d-e** and **S7h-i**). Altogether, these results indicate that the principal neurons of dKAT3-KOs lose their neuronal identity, but do not die, dedifferentiate or transdifferentiate to other cell types. Instead, these cells seem to be trapped in a non-functional interstate deadlock.

### CBP and p300 bind to the same regulatory regions

The evidence above indicates that KAT3 proteins play a redundant role preserving neuronal identity. To explore the basis of such redundancy, we next mapped the occupancy of hippocampal chromatin by CBP and p300. Chromatin immunoprecipitation followed by whole-genome sequencing (ChIP-seq) using antibodies that differentiate between the two paralogous proteins (**Fig. S1b** and **S8a**) retrieved 37,359 peaks in the chromatin of wild type mice (**Fig. 4a, S8b** and **Table S4**). After correcting for the different efficiencies of the two antibodies, we detected an almost complete overlap between the CBP and p300 peaks throughout the whole genome (**Fig. 4b** and **S8c**). This finding, supported for the combined analysis of genome occupancy by CBP and p300 in wild type animals and single and double ifKOs, reveals for the first time that these two essential epigenetic enzymes occupy basically the same sites in neuronal chromatin, and underscores the large functional redundancy of KAT3 proteins.

**Figure 4.**
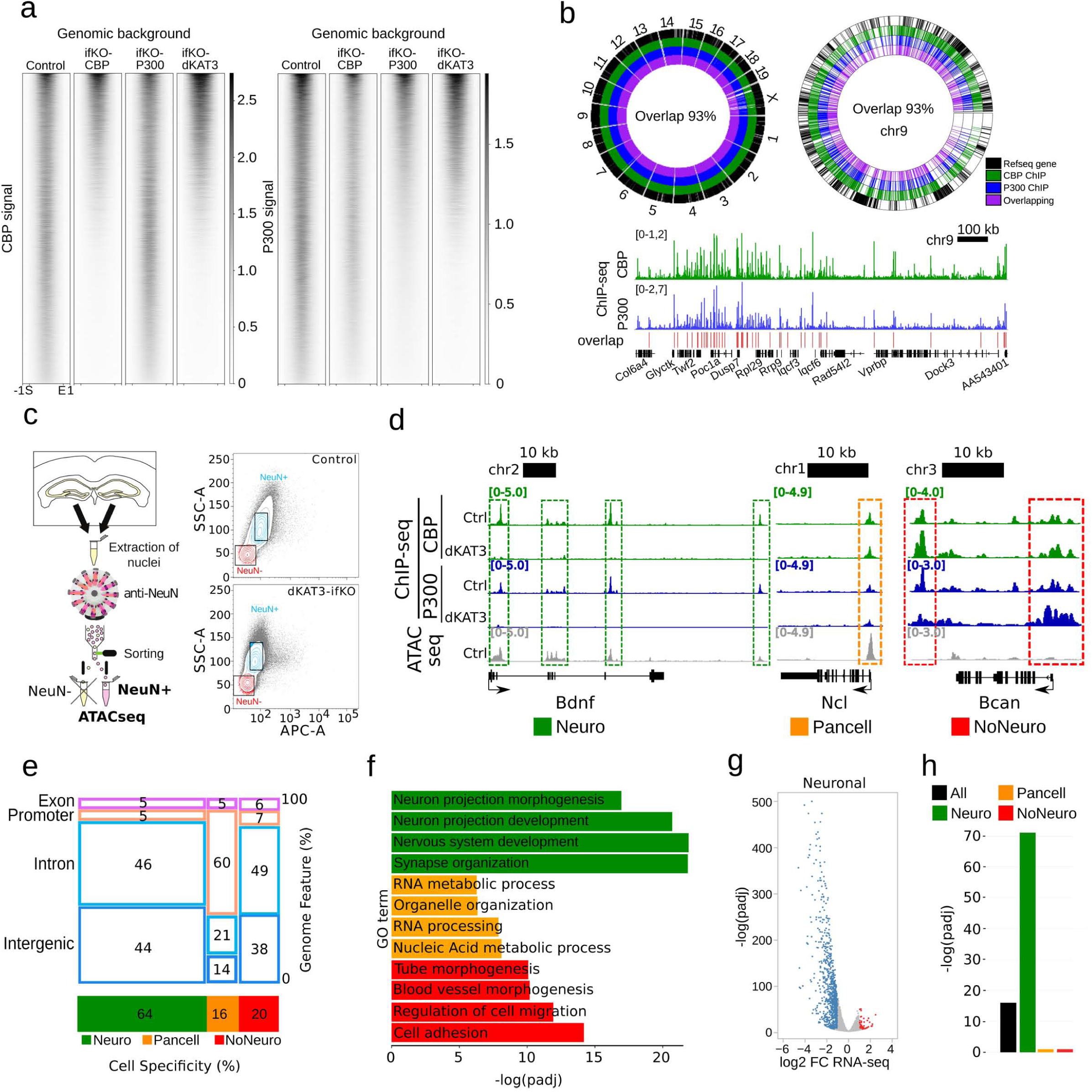
CBP and p300 bind to the same genomic sites. **a.** Heat maps showing the control CBP/P300 KAT3 ChIP-seq peaks and the signal in the corresponding locations of CBP-, p300- and KAT3-ifKO. Intensity ranges from strong (black) to weak (white). S: peak start, E: peak end, +/− 1kb. **b.** Circos plots of the entire genome (left) and chromosome 9 (right). Below: a snapshot of a gene-rich region in chromosome 9. Colors indicate CBP and p300 binding in hippocampal chromatin and their overlap. Refseq genes in black **c.** Left: Scheme of the FANS/ATAC-seq experiment. Right: Flow cytometry sorting plots for control and dKAT3-fKO samples. Boxes indicate the gates used for sorting NeuN+ nuclei (blue). APC-A is the signal of anti-NeuN staining. **d.** Snapshot illustrating the classification of KAT3 peaks in neuronal (green), non-neuronal (red) and pancellular (orange). Representative peaks classified as neuronal, non-neuronal and pancellular are marked with a green, red and orange dashed rectangle, respectively. **e.** Classification of KAT3 peaks according to cell specificity (neuronal, pancellular and non-neuronal) and genomic feature (promoter, exon, intron, intergenic). The numbers within each sector represent percentages. **f.** GO enrichment analysis performed on the gene sets associated with neuronal (green), non-neuronal (red), and pancellular (orange) KAT3 peaks. **g**. Volcano plots of RNA-seq analysis for the subset of genes classified as neuronal. Red: upregulated genes; blue: downregulated genes. **h.** BETA analysis of the association between all, neuronal, non-neuronal and pancellular KAT3 peaks, and transcriptome changes. Neuronal peaks show the strongest association with gene downregulation.

As both proteins are ubiquitously expressed yet gene ablation only takes place in excitatory neurons, the signal detected in the hippocampal chromatin of dKAT3-ifKOs must correspond to CBP/p300 binding in other cell types. We combined this information with ATAC-seq (a technique that investigates chromatin occupancy and requires much lower input than ChIP-seq) in sorted NeuN^+^ hippocampal nuclei ^25^ to discriminate between neuronal-specific, non-neuronal-cell-specific and “pancellular” KAT3 binding (**Fig. 4c-d, S9a** and **Table S4**). Intriguingly, pancellular KAT3 peaks are found in the promoter of genes involved in basic cellular functions, such as RNA processing and metabolism, while cell-type-specific peaks (neuronal and non-neuronal) primarily locate at introns and intergenic regions with enhancer features (**Fig. S9b**) and associate with cell-type-specific processes (**Fig. 4e-f** and **Table S5**). We used binding and expression target analysis (BETA) ^26^ to integrate ChIP-seq and RNA-seq data (**Fig. 4g**), and found that the loss of neuronal KAT3 binding (∼14,000 peaks) is an excellent predictor of transcriptional downregulation (p = 3.3×10^−72^, **Fig. 4h**). Up to 74% of the downregulated genes in dKAT3-ifKOs are linked to the loss of KAT3s at proximal regulatory regions. Comparison of the ATAC-seq accessibility profiles of control and dKAT3-ifKO neuronal nuclei retrieved more than 6,000 differentially accessible regions (DARs) (**Table S6**). Most of these DARs displayed reduced accessibility in dKAT3-ifKO neurons (**Fig. S9c**), were located at enhancers (**Fig. S9d**) and coincided with the downregulation of the proximal gene (**Fig. S9e**).

### KAT3 proteins control the acetylation levels of neuron-specific enhancers

The acetylation of histone H3 at lysine 27 (H3K27ac) is a likely mechanism for mediating the role of KAT3 proteins in maintaining neuronal identity. This histone modification is enriched in active enhancers and its levels correlate with tissue specification ^27^. In agreement with a recent acetylome analysis ^11^, immunostaining against specific lysine residues in the histone tails revealed their variable dependence on CBP/p300 (**Fig. S10a**) as well as the particular sensitivity of H3K27ac to the loss of CBP and p300 (**Fig. 5a**). The dramatic reduction in H3K27ac is not accompanied by an increase in the signal or changes in the distribution of H3K27me3 (**Fig. S10b**). H3K9me3, a histone modification associated with heterochromatin, also appears unaffected (**Fig. S10c**).

**Figure 5.**
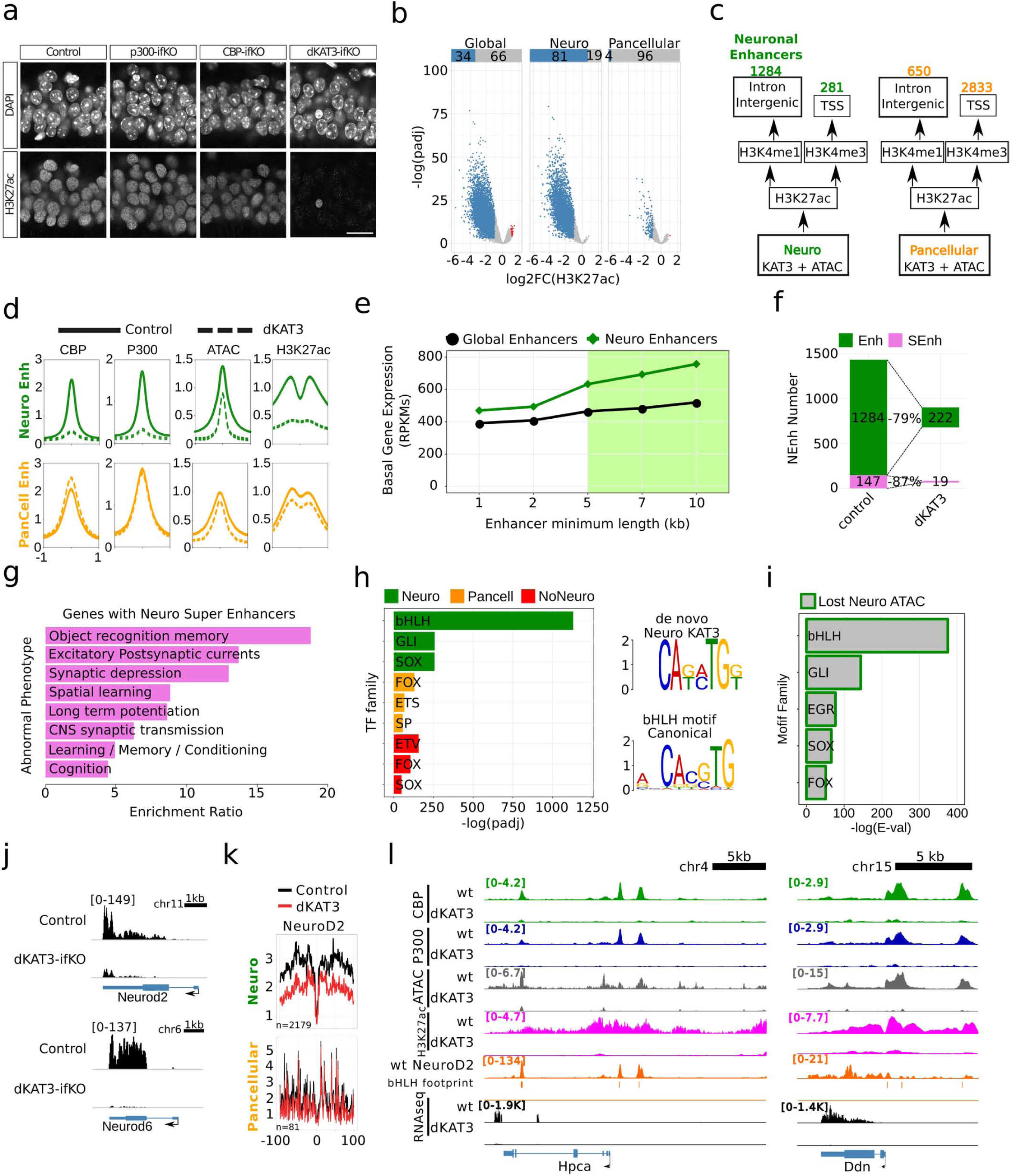
H3K27ac is strongly decreased in neuro-specific locations and correlates with gene downregulation. **a**. Immunostaining against H3K27ac in the CA1 subfield of single and dKAT3 ifKOs. Scale: 10 µm. **b.** Genome-wide analysis of H3K27ac changes in dKAT3-ifKOs. Values are shown for all peaks and split between neuronal and pancellular peaks. **c.** Classification of regulatory regions categorized by genomic features. **d.** Metaplots of ATAC-seq, KAT3 and H3K27ac ChIP-seq signals in neuronal and pancellular enhancers in controls and dKAT3-ifKOs. Plots are centered in the peak center and expanded +/− 1 kb **e**. Correlation between length of the enhancer and expression level of the proximal gene. Note that expression raises for enhancers longer than 5 kb. **f.** Number of downregulated genes that contain enhancers or super-enhancers in control mice and the percentage that lose acetylation in dKAT3-ifKOs. **g.** Barplot showing the top 10 enriched categories from the *Phenotype* analysis for genes harboring neuronal enhancers performed with the WEB-based GEne SeT AnaLysis Toolkit. **h.** TFBS analysis of neuronal (green), non-neuronal (red) and pancellular (orange) KAT3 peaks. Each motif family name is a user-curated approximation to the results provided by the MEME-suite algorithm. The most enriched *de novo*-identified binding motif in neuronal KAT3 (upper right motif) is very similar to the NeuroD2 motif (bottom right). **i.** TFBS analysis of neuronal regions with a reduced ATAC-seq signal in dKAT3-ifKOs. **j.** RNA-seq profiles for *Neurod2* and *NeuroD6*. Scale: 1 kb. **k.** Digital footprinting of NeuroD2 at neuronal and pancellular KAT3-bound regions. Values correspond to normalized Tn5 insertions. **l.** Representative snapshots of KAT3, ATAC and H3K27ac depletion with bHLH footprint and NeuroD2 overlaps, at two genes that are strongly downregulated in dKAT3-ifKOs.

We next investigated the genomic distribution of H3K27ac in the hippocampal chromatin of dKAT3-ifKOs and control littermates. We retrieved 37,732 H3K27ac-enriched regions (**Fig. 5b**) that largely overlap with KAT3 peaks (**Fig. S10d**). More than one third of these H3K27ac peaks were strongly reduced in dKAT3-ifKOs (**Table S7**), particularly those that overlap with neuronal KAT3 binding (∼80%). In contrast, less than 5% of pancellular and non-neuronal H3K27ac-enriched regions were affected (**Fig. 5b**). To explore in greater detail these differences, we classified neuronal KAT3 peaks into promoters and enhancers based on their location and H3K4me1/me3 content (**Fig. 5c**). In dKAT3-ifKOs, KAT3 binding, H3K27ac and ATAC-seq signals were all strongly reduced in neuronal enhancers (**Fig. 5d** and **S10e**), while the promoters associated with pancellular-KAT3 peaks were spared, indicating that other KATs maintain the acetylation of these loci. Consistent with this view, the acetylation of H3K9,14 that decorates promoters only showed a modest reduction in dKAT3-ifKOs (**Fig. S10f**), indicating that other KATs are responsible for maintaining this form of acetylation at promoters. We also observed a strong correlation between the loss of H3K27ac and transcript downregulation. Of the 1,376 downregulated genes in dKAT3-ifKOs, 78% showed a strong reduction in H3K27ac. Reciprocally, 74% of the genes with reduced acetylation were neuronal genes severely downregulated in dKAT3-ifKOs.

Super-enhancers constitute a special type of regulatory region that results from a cluster of several enhancers bound by master regulators and the Mediator complex and plays a critical role in controlling cell identity ^28^. Their conspicuous features include the association with highly transcribed genes, broad domains of H3K27 acetylation, and a high density of TF binding sites. To identify putative neural super-enhancers, we fused together enhancers that are closer than 5 kb from each other (**Table S8**). The retrieved super-enhancers were associated with highly expressed genes in hippocampal neurons that are downregulated in dKAT3-ifKOs (**Fig. 5e**). Furthermore, most of these super-enhancers-associated genes were strongly hypoacetylated in dKAT3-ifKOs (**Fig. 5f**) and encode *bona fide* neuronal regulators related with synaptic transmission and neuronal plasticity (**Fig. 5g**). Neuronal fate also relies on changes in chromatin architecture during neural differentiation ^29^. In agreement with this view, we found that the loss of chromatin accessibility in dKAT3-ifKOs overlapped with the establishment of cell type-specific enhancer-promoter contacts during neural differentiation, which is concomitant with the activation of neuron-specific transcription (**Fig. S10g-h**). Together, these results show that KAT3 proteins control the status of neuron-specific genes by regulating lysine acetylation levels and chromatin interactions at enhancers and super-enhancers.

### bHLH TFs drive KAT3 binding to neuron-specific genes and enhancers

Since KAT3 proteins do not directly bind to DNA, we next asked which proteins are responsible for recruiting CBP and p300 to these neuron-specific regulatory regions. Motif prediction analysis of KAT3 binding peaks revealed remarkable differences between pancellular and neuronal-specific regions. Pancellular KAT3 binding is associated with general TFs such as Sp, Fox and Ets, all of which have been reported to bind to CBP or p300 ^10^ (**Fig. 5h**). In contrast, neuronal KAT3 peaks presented a very prominent enrichment (E-value = 1.0_x_10^−1223^) for the DNA binding motif of basic helix-loop-helix (bHLH) proteins (**Fig. 5h**). The regions losing occupancy in dKAT3-ifKOs were also highly enriched for bHLH-recognized motifs (**Fig. 5i** and **S11a**).

Among the bHLH proteins there are pro-neural TFs critically involved in neuronal development ^30, 31^ (e.g., *Ascl1*, *Neurod1-6*, *Neurogenin1-3* and *Atoh1*). Our differential expression analysis retrieved 20 bHLH encoding genes that are downregulated in the hippocampus of dKAT3-ifKOs (**Fig. S11b** and **Table S2**). In fact, the *NeuroD2* and *NeuroD6* genes, which encode TFs with putative terminal selector activity ^5, 32^, were among the most downregulated genes in the hippocampi of dKAT3-ifKOs (**Fig. 5j**). These results suggest that the change in accessibility is a likely consequence of the loss of both the KAT3 proteins and the recruiting TFs. To directly assess the occupancy of these sites before and after the ablation of KAT3s, we analyzed their digital footprints in DARs and detected robust differences for pro-neural bHLH TFs (**Fig. 5k** and **S11c**), confirming that some changes in accessibility reflect the reduced binding of one of several member of this family. This result is consistent with the large overlap between the bHLH footprints detected in the ATAC-seq profiles and the Neurod2 ChIP-seq data ^33^ (**Fig. S11d**). Overall, these experiments indicate that CBP and p300 interact with bHLH proneural TFs in neuronal-specific genomic locations to maintain neuron-specific transcription (**Fig. 5l** and **S11e** show some representative loci).

### Other cell types also require KAT3 protein for maintaining their identity

Previous evidence indicates that KAT3 proteins play a critical role in the differentiation of other cell types and the establishment of cell type-specific transcription ^34, 35, 36, 37^. To investigate if cell fate maintenance in other cell types also requires the KAT3 proteins, we next examined astrocytes derived from dKAT3-floxed mice (**Fig. 6a**). We prepared astrocyte cultures from the hippocampi of *Crebbp^f/f^::Ep300^f/f^* embryos and infected them with a Cre-recombinase-expressing lentivirus. Similar to our results in neurons, primary cultures of cortical astrocytes missing both KAT3 proteins lose both the expression of glial genes such as *Gfap* (**Fig. 6b-c**) and their characteristic morphology (**Fig. 6d**), indicating that KAT3 proteins are also responsible for identity maintenance in other cell types.

**Figure 6.**
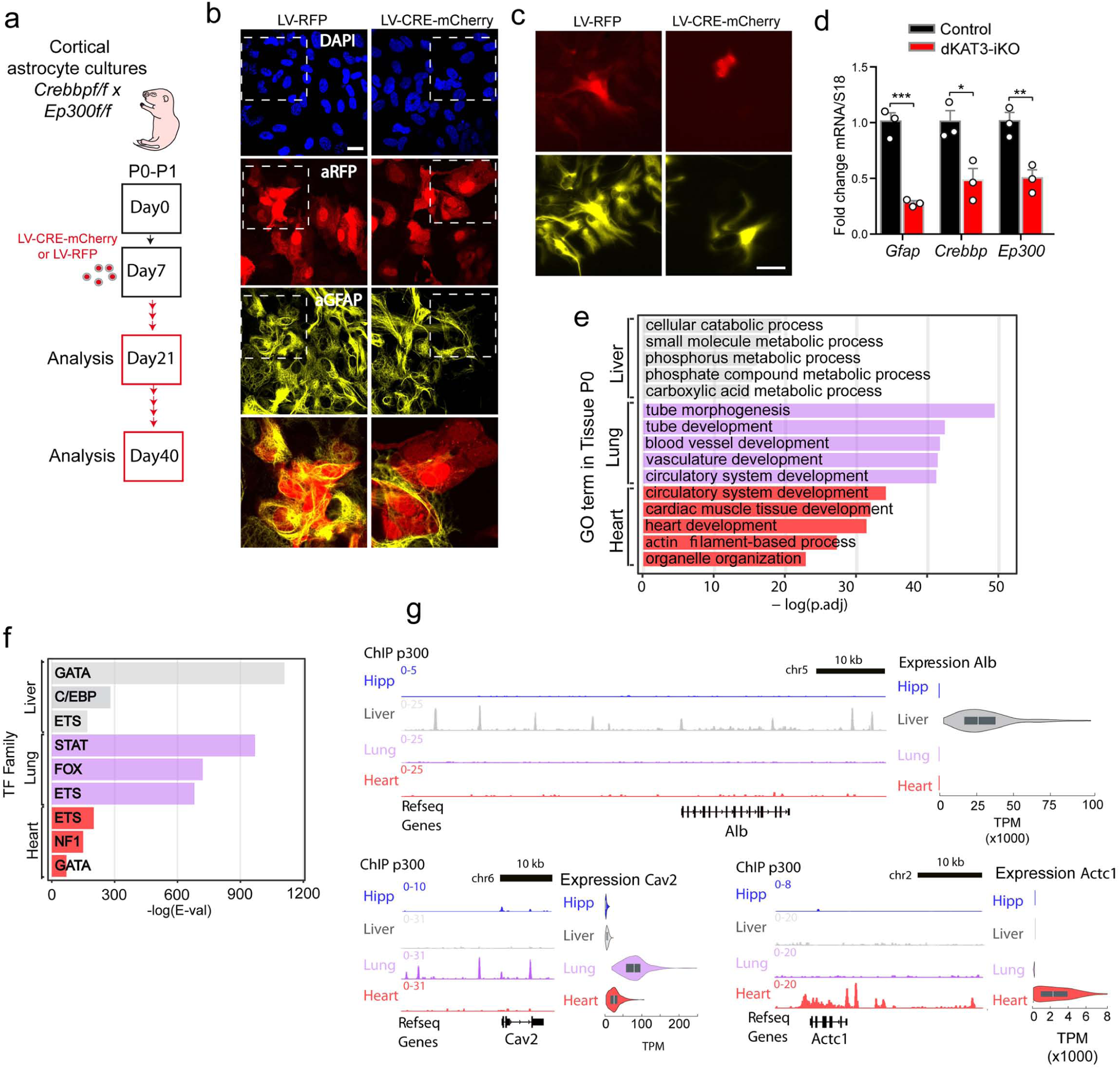
CBP and p300 are needed to maintain the fate of other cellular types. **a.** Generation of dKAT3-KO astrocytes. Cortical astrocytes from P0 *Crebbp*^f/f^::*Ep300*^f/f^ pups were infected with a cre recombinase-expressing LV. **b.** Cultured dKAT3^f/f^ astrocytes show a downregulation of astrocyte marker GFAP 2 weeks after infection with a cre recombinase expressing LV. Scale bar: 20 µm. **c.** RT-PCR quantification of the *Gfap* (GFAP), *Crebbp* (CBP) and *Ep300* (p300) transcript levels in cultured dKAT3^f/f^ astrocytes. **d.** The loss of astrocyte morphology is more evident 4 weeks after infection. Scale bar: 50 µm. **e.** Top five most enriched categories identified by GO analysis on the gene sets associated with p300 binding in liver, lung and heart of P0 mice. Categories correspond to expected functions for each organ. **f.** TFBS analysis of p300-bound putative enhancers (intergenic and intronic peaks) in chromatin of liver, lung and heart tissue from P0 mice. Motif family names are taken from the most prominent results provided by the MEME-ChIP algorithm. In the case of lung and heart we detected large enrichments for specific TFs but with no clear foremost candidate as in hippocampus and liver, likely reflecting the greater cellular heterogeneity of these tissues. **g.** Representative snapshots of p300 ChIP-seq in hippocampus (this study), and liver, lung and heart (ENCODE) at three representative tissue-specific genes. Gene expression in Transcripts per Million reads (TPM) for each tissue was obtained from the Genotype-Tissue Expression (GTEx) project (www.gtexportal.org) and it is shown as violin and box plots adjacent to each track. *Alb*, *Cav2* and *Actc1* encode for Albumin, Caveolin 2 and Actin alpha cardiac muscle 1, respectively.

To examine this possibility in cells from other lineages, we investigated the profiles of p300 binding in different tissues available at ENCODE (there are no tissue-specific profiles for CBP). We found that, similarly to our observations in brain tissue, p300 peaks were associated with genesets highly enriched in tissue-specific functions (**Fig. 6e**). Furthermore, in each tissue, p300-bound regulatory regions were enriched for distinct binding motifs (**Fig. 6f**). Profuse and tissue-specific p300 binding was observed into and upstream of loci presenting tissue-specific transcription (**Fig. 6g**). The most remarkable example is the liver, in which we detected a very high enrichment for GATA TFs similar to the one observed for bHLH TFs in excitatory neurons. TFs belonging to this family are essential during liver development and are still expressed in mature tissue ^38^, indicating that they may work as terminal selector for hepatocytes.

### The molecular scaffold and KAT activities of CBP are both required to re-establish neuronal transcription

To explore the specific contribution of the scaffolding and KAT activities of KAT3 proteins to the loss of cell type-specific transcription, we turned to PNCs. Importantly, transfection of a heterologous full-length CBP in neurons that have loss the endogenous expression of both CBP and p300 (**Fig 7a-b**) fully prevented the rapid loss of expression and neuronal markers (**Fig 7c-d**). We next examined whether the expression of the N-terminus or the C-terminus (bearing the KAT domain) halves of CBP, as well as the full-length reconstituted protein (**Fig. 7e-f**) could also rescue the transcriptional impairment. The specifically assess the contribution of KAT activity, we also assessed a variant of the C-terminus half of CBP (referred to as KATmut) bearing the R1378P mutation linked to RSTS ^39^. We found that only the reconstruction of full-length CBP with an intact KAT domain prevented the downregulation of neuronal markers (**Fig. 7g-h**). These results indicate that both activities of CBP, as KAT and molecular scaffold, are necessary to preserve locus activity. We also investigated whether heterologous NeuroD2 expression exerted the same protection. This bHLH TF is highly expressed in mature excitatory neurons, regulates its own expression and holds the features of a neuronal terminal selector ^5^. Furthermore, it is strongly downregulated in dKAT3-ifKOs neurons (**Fig. 5j** and **S11b**), and its occupancy profile in cortical chromatin ^33^ shows a large overlap with DARs in dKAT3-ifKOs (**Fig. S12a-b**). However, conversely to CBP, NeuroD2 alone was not sufficient to re-establish neuronal-specific transcription in dKAT3-KO PNCs (**Fig. S12c**).

**Figure 7.**
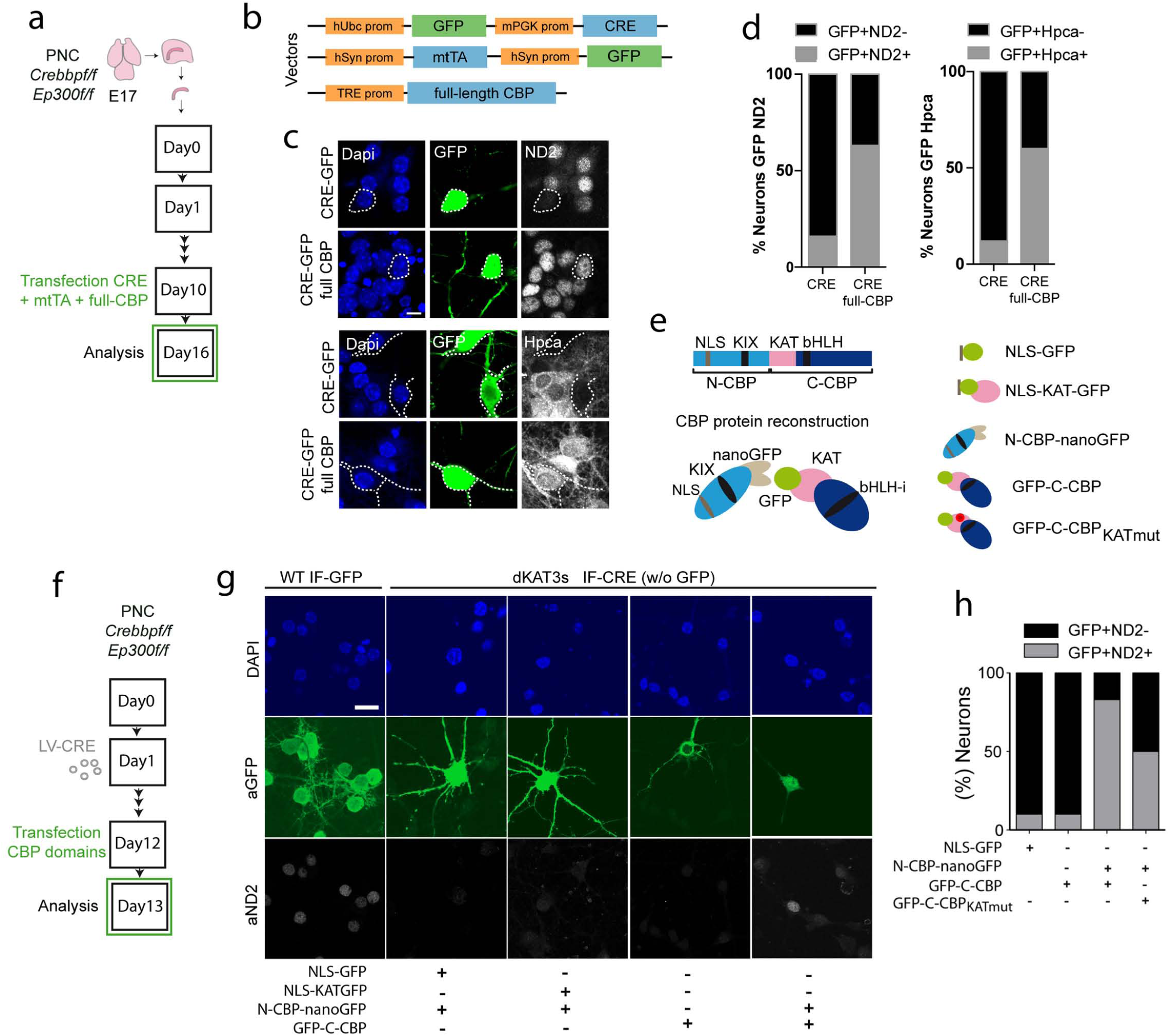
Full-length CBP is required to restore neuronal-specific transcription. **a.** Hippocampal PNCs from E17 *Crebbp*^f/f^::*Ep300*^f/f^ embryos were co-transfected with constructs that drive the expression of the Cre recombinase and full-length CBP. Plasmid combination to express recombinant CBP simultaneously to endogenous CBP and p300 ablation. **c**. Representative images of NeuroD2 and hippocalcin staining in PNC transfected with the constructs shown in **panel 7b**. Note the reduced expression in the GFP+ cells in the absence of heterologous full-length CBP. **d**. Quantification of the percentage of NeuroD2-positive or -negative cells and hippocalcin-positive or -negative cells among all GFP-positive cells. **e**. Scheme of the CBP fragments used for rescuing NeuroD2 expression (see Materials and Methods for additional details). NLS: Nuclear localization domain; KAT: acetyltransferase domain; KIX: kinase-inducible domain interacting domain; bHLH-i: region of interaction with bHLH transcription factors. **f.** Hippocampal PNCs from E17 *Crebbp*^f/f^::*Ep300*^f/f^ embryos were transfected with constructs that drive the expression of the Cre recombinase and the different CBP fragments and the KAT domain shown in **panel 7e**. **g**. Representative images of NeuroD2 staining in PNC infected with LV-CREw/oGFP and transfected with the different domains of CBP shown in **panel 7e**. LV-GFP was used as a control of the baseline NeuroD2 level (2 independent PNCs). Scale: 20 µm. **h**. Quantification of the percentage of NeuroD2-positive or -negative cells among all GFP-positive neurons.

### Locus-specific epi-editing rescue transcriptional impairments

Next, we examined whether increasing lysine acetylation is sufficient to rescue the transcriptional deficit. To this end, we took advantage of recently developed tools for epi-editing based on the expression of an inactive Cas9 enzyme (dCas9) fused to the KAT domain of p300 (dCas9-KAT) ^40^. This system enables targeting specific genomic regions for p300-dependent acetylation using adequate guide RNAs (gRNA). We selected *Neurod2* as the target gene because it is both severely downregulated and H3K27-deacetylated in dKAT3-ifKOs (**Fig. 8a**). The infection of PNCs with lentiviruses that express dCas9-KAT and a gRNA that recruits this chimeric protein to the most proximal KAT3 peak of *Neurod2* (**Fig. 8a-b**) prevented the downregulation of NeuroD2 that was observed in dKAT3-KO neurons (**Fig. 8c-d**). Moreover, co-transfection of plasmids carrying dCas9-KAT and the Neurod2 gRNA in dKAT3-KO cells that had already lost Neurod2 expression also caused a significant recovery of the expression of this gene (**Fig. 8e-g**). These experiments demonstrate that the increase of KAT activity at the locus preserves and can even restitute its functionality, suggesting that histone/lysine deacetylation at regulatory regions is the main cause for the downregulation of neuronal genes after KAT3 loss.

**Figure 8.**
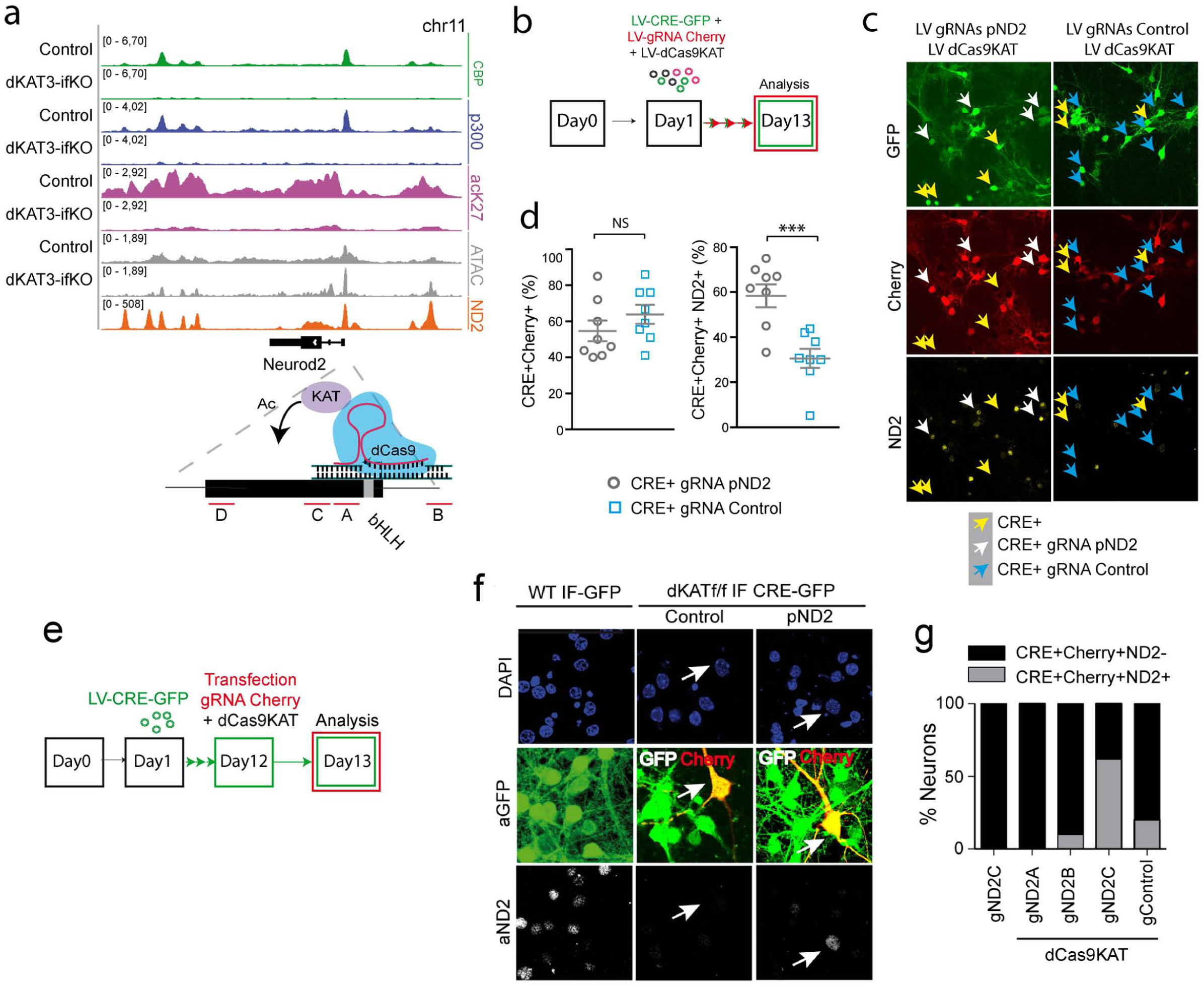
Locus-specific acetylation restores NeuroD2 transcription. **a**. Snap view of the Neurod2 locus. The profiles for CBP and p300 binding, H3K27 acetylation and ATAC-seq signal are shown. A scheme representing the strategy used to drive p300 KAT activity to a specific genomic locus using dCas9-KAT system and the location of the sequences at the *Neurod2* promoter targeted by the gRNAs (red lines, gND2 A-D) are also shown. Note that the target regions overlap with the bHLH sites at the NeuroD2 promoter retrieved in ^33^ (bottom orange track). **b.** Scheme of the co-infection of LVs expressing cre recombinase, dCas9-KAT and the Neurod2 gRNA to prevent Neurod2 loss in dKAT3-KO neurons. **c.** Representative image of NeuroD2 protein levels in the cells co-infected with LV-CRE-GFP and LV-gRNA-mCherry specific for *Neurod2* locus (white arrows), the specificity of gRNAs was demonstrated using targeted gRNAs specific for *Hpca locus* (Hip; blue arrows). As a comparison, cells infected with LV-CRE-GFP alone (yellow arrows) show a strongly diminished NeuroD2 levels as well. Scale bar: 50 µm. **d.** Quantification of different cell subpopulations observed in the experiment shown in panel **8c**. **e.** Scheme of rescue experiment with plasmids carrying dCas9-KAT and the Neurod2 gRNA in dKAT3-KO cells that had already lost Neurod2 expression. **f.** Representative image of NeuroD2 expression after transfection with dCas9-KAT and gND2-C targeting the NeuroD2 promoter. Arrows indicate LV-CRE-GFP infected cells transfected with the gRNA-carrying vector. A gRNA targeting the hippocalcin promoter was used as a specificity control. Scale: 20 µm **g.** Quantification of the percentage of transfected cells showing normal NeuroD2 expression after transfection with gND2-C alone and dCas9-KAT co-transfected with gND2-C, gND2-A, gND2-B or gRNA control independently (experiments in three independent PNCs).

## Discussion

### From gate-to fate-keepers

The differentiation of neuronal lineages during brain development requires the participation of TFs and chromatin-modifying enzymes such as CBP and p300 ^6, 12, 41, 42, 43, 44, 45, 46^. These proteins are often referred to as *gatekeepers* of cell fate. Here, we used an inducible knockout system to demonstrate that KAT3 proteins are jointly required in excitatory neurons to preserve their postmitotic neuronal identity, thereby acting as *fate-keepers*. This evidence indicates that the epigenetic landscape of neuronal genes is not self-sufficient and requires the active and continuous presence of KAT3 enzymes, which opens up the possibility of using controlled KAT3 inhibition for dedifferentiation and reprograming in cell therapy strategies. Supporting this view, a recent study showed that treatment with competitive inhibitors targeting the bromodomains of CBP and p300 enhanced the reprogramming of human fibroblasts into iPSCs, while KAT inhibition prevented iPSC formation ^47^ (which is consistent with our results demonstrating that dKAT3-KO fail to express stem cell markers).

Strikingly, the loss of neuronal identity did not lead to apoptosis or other forms of neuronal death even weeks after KAT3 elimination both in culture and *in vivo*. This may be result of the inability of dKAT3-KO cells to trigger the programed cell death program. As shown in **Fig. S4b**, numerous genes involved in neuronal apoptosis and death are strongly downregulated in dKAT3-KO, including important initiators of neuronal apoptosis such as *Hrk* ^48, 49^. This gene is highly expressed in neurons and as many other neuronal genes present a dramatic downregulation that is accompanied by the loss of CBP/p300 binding and the H3K27 deacetylation of the locus (**Fig. S11e, left**). Together these results suggest that KAT3 proteins are essential to safeguard cell identity but also to activate any alternative cell fate, including stemness and programmed cell death.

### Molecular mechanism of cell fate maintenance

Thanks to their ability to interact with cell type-specific TFs, KAT3 proteins are recruited to specific genomic locations where they act as a ‘‘acetyl-spray’’ targeting accessible lysine residues on proximal histones and non-histone proteins ^11^ to support active transcription ^12^. Interestingly, a single *Crebbp* or *Ep300* functional allele is sufficient to sustain neuronal identity, which underscores the robustness of the epigenetic mechanisms involved in cell type maintenance. Robustness and redundancy are even greater in ubiquitous and highly expressed genes for which we detected neither downregulation nor severe hypoacetylation in dKAT3-ifKOs. This indicates that other KATs must maintain acetylation levels at these loci. Consistent with this hypothesis, our RNA-seq screen shows that other KATs such as Gcn5/KAT2A, Tip60/KAT5 and MOF/KAT8, which are expressed in neurons and localize to promoters ^50, 51^, are not downregulated in dKAT3-ifKO neurons (**Fig. S11e, right** shows the profile of *Kat5*, a similar pattern is observed in *Kat2A* and *Kat8*).

Our experiments show that the recruitment of KAT3 *proteins* to cell type-specific enhancers in excitatory hippocampal neurons is likely mediated by bHLH proteins from the NeuroD family. Various bHLH TFs are differentially expressed throughout neuronal proliferation and specification. For instance, Hes1 and Hes5 promote NSC/NPC renewal and inhibit specification, but are replaced by Ascl1 or Neurog2 to trigger the differentiation to cortical neurons ^52–54^. These two TFs are followed or accompanied by other factors like NeuroD1 and other family member, such as NeuroD2 or NeuroD6, whose expression is maintained throughout the lifetime of the neuron ^55^. Although we cannot be certain which specific bHLH TF or set of TFs actually recruit CBP and p300 in adult neurons, the two KATs are known to interact with multiple bHLH TFs including MyoD ^56^, Ascl1 ^41^, Neurogenins ^41, 42, 57^, Twist ^58^, and NeuroD2 ^59, 60^. Since bHLH proteins form dimers ^54, 61^, and binding to a dimerized TF has recently been shown as essential for the control of KAT3 activity ^62^, these TFs represent suitable candidates to regulate KAT3 function and drive chromatin acetylation at different stages of neurodevelopment. The fact that the neural tube phenotype of *Crebbp^−/−^* mice mimics the phenotype of *Twist* null knockouts is consistent with this hypothesis ^13, 63^. The loss of neuronal identity that is observed in adult neurons of dKAT-ifKOs is unlikely to be a direct consequence of the downregulation of pro-neural TFs because this phenotype was not observed after elimination of *Neurod1*, *Neurod2*, *Ascl1* or other bHLH genes in neurons ^55, 64, 65^, nor was this phenotype rescued by NeuroD2 overexpression. It is also unlikely to be the direct consequence of a general loss of the enhancer’s building factors because our differential gene expression analysis indicates that, besides the two KAT3 proteins, the components of the enhanceosome ^66^ are not downregulated in dKAT3-ifKOs. Instead, our experiments show that identity loss is strongly associated with chromatin hypoacetylation and that it can be avoided by targeted KAT activity. These results suggest that the main role of terminal selectors might be to recruit the KAT3 enzymes for maintaining the acetylated status of cell type-specific enhancers in differentiated cells. These findings have broad clinical implications because impaired CBP/p300 function, histone hypoacetylation and the loss or attenuation of the epigenetic profiles underlying cell fate are features of several neurological disorders, including Alzheimer’s and Huntington’s diseases and aging-related senescence ^67, 68, 69^.

### A conserved role in cell fate maintenance

Importantly, the role of KAT3 proteins as fate-keepers is not restricted to neurons. Our experiments in astrocytes, analysis of KAT3 peaks in different tissues and previous observations in other cell types demonstrating that the combined loss of CBP and p300 causes impaired cell type-specific transcription ^34, 35, 36, 37^, all support this view. What may differ between cell types is the specific set of TFs that recruit the KAT3 proteins to cell type-specific loci (the bHLH and GATA family of TFs in the tissues explored here). Further studies should be aimed at identifying the cell type-specific partners and targets governing tissue specification and signaling.

The role of KAT3 proteins in preserving cell type-specific gene statuses can be particularly relevant for neurons given their tremendous diversity, elaborate connectivity patterns and long lifespan. The critical importance of KAT3s for brain function may explain the duplication of the ancestral KAT3 gene in the first vertebrates coinciding with the emergence of a neural crest and cephalization ^12, 70^. Although the two KAT3 proteins have evolved some individual functions in postmitotic cells (e.g., forebrain restricted CBP knockouts show phenotypes related to cognitive dysfunction ^67^ and p300 seems to play a prominent role in muscle biology ^71^), our study demonstrates that they still share a joint and more essential role preserving epigenetic identity.

## Supporting information

Experimental procedures and Supplementary Figures

## Acknowledgments

We thank P. Arlotta, O. Hobert, N. Flames, E. Herrera, M.A. Nieto, A. Rada-Iglesias and J.V. Sanchez-Mut for critical reading of the manuscript. We thank A. Caler, N. Cascales-Picó, M. Llinares, A. Medrano-Fernández and S. Rivero for their assistance in specific experiments and V. Makarov for the ICAofLFPs MatLab package. M.L. is recipient of a *Santiago Grisolia* fellowship given by the Generalitat Valenciana, J.M.C. is recipient of a fellowship from the Spanish Ministry of Education, Culture and Sport (MECD), J.F-A. and C.M.N. are recipients of fellowships from the Spanish Ministry of Science and Innovation (MICINN). The ultrastructure research was supported by the Polish National Science Center grant UMO-2014/15/N/NZ3/04468 and by the European Regional Development Fund POIG 01.01.02-00-008/08. J.L-A. research is supported by grants RYC-2015-18056 and RTI2018-102260-B-I00 from MICINN co-financed by ERDF. A.B. research is supported by grants SAF2017-87928-R, PCIN-2015-192-C02-01 and SEV-2017-0723 from MICINN co-financed by ERDF, PROMETEO/2016/026 from the Generalitat Valenciana, and RGP0039/2017 from the Human Frontiers Science Program Organization (HFSPO). The Instituto de Neurociencias is a “Centre of Excellence Severo Ochoa”.

## Author contributions

Conceptualization, M.L, R.M-V., B.dB. and A.B.; Methodology, M.L, B.dB., J.F-A., C.M.N., R.O., A.A.S., J.M.C.; Software, R.M-V., and A.M-G.; Investigation, M.L., B.dB., and J.M-R., Data Curation and Visualization, R.M-V., A.M-G., M.L.; Writing – Original Draft, A.B. and M.L.; Supervision, A.B. and J.L-A., S.C., G.M.W.; Funding Acquisition, A.B.

