## Supplementary material for "Neuronal identity is maintained in the adult brain through KAT3-dependent enhancer acetylation": Experimental procedures and Supplementary Figures

**Inventory of Supplementary materials**


- Experimental Procedures (pages 3-20)
- Supplementary References (pages 21-22)
- Supplementary figures S1-S12 and supplementary legends (pages 23-47)
- Supplementary Tables:
  - **Table S1 related to Figure 1.** Results of SHIRPA screen in single and double KAT3 ifKOs (page 48).
  - **Table S2 related to Figure 2.** 3 sheets: (i) RNA-seq sample description; (ii) Differentially gene expression analyses for all genes; (iii) deregulated genes in the dKAT3-ifKO RNA-seq (see attached Excel file).
  - **Table S3 related to Figure 3.** Top population markers and results from differential expression analysis to identify population-enriched genes (see attached Excel file).
  - **Table S4 related to Figure 4.** 5 sheets: (i) CBP and P300 ChIP-seq sample description; (ii) bed file for all KAT3 peaks; (iii) Differential KAT3 binding in dKAT3-ifKO vs control for neuronal KAT3 peaks; (iv) Differential KAT3 binding in dKAT3-ifKO vs control for pancellular KAT3 peaks; (v) Differential KAT3 binding in dKAT3-ifKO vs control for non-neuronal KAT3 peaks (see attached Excel file).
  - **Table S5 related to Figure 4.** 3 sheets: (i) GO Enrichment analysis for genes annotated to closest neuronal KAT3 peaks; (ii) GO Enrichment analysis for genes annotated to closest pancellular KAT3 peaks; (iii) GO Enrichment analysis for genes annotated to closest non-neuronal KAT3 peaks (see attached Excel file).

- **Table S6 related to Figure 4.** 4 sheets: (i) ATAC-seq sample description; (ii) Global Differential Accessibility in dKAT3-ifKO vs control for KAT3 peaks; (iii) Differential Accessibility in dKAT3-ifKO vs control for neuronal KAT3 peaks; (iv) Differential Accessibility in dKAT3-ifKO vs control for pancellular KAT3 peaks (see attached Excel file).
- **Table S7 related to Figure 5.** 5 sheets: (i) H3K27ac ChIP-seq sample description; (ii) Differential H3K27ac amount for all peaks; (iii) Differential H3K27ac amount in dKAT3-ifKO vs control for neuronal peaks; (iv) Differential H3K27ac amount in dKAT3-ifKO vs control for pancelular peaks; (v) Differential H3K27ac amount in dKAT3-ifKO vs control for non-neuronal peaks (see attached Excel file).
- **Table S8 related to Figure 5.** 3 sheets: (i) list of all detected enhancers (Enh) and super enhancers (SEnh); (ii) list of neuronal specific enhancers (Enh) and super enhancers (SEnh) detected in controls; (iii) list of neuronal specific enhancers (Enh) and super enhancers (SEnh) lost in dKAT3-ifKO (see attached Excel file).

- Supplementary Movie:
  - **Movie M1 related to Figure 1.** Rapid emergence of severe neurological phenotypes in dKAT3-ifKOs (MP4 file).

#### Experimental procedures

*Animals and treatments*

Previously described CaMKIIα-creERT2 [1], *Ep300*$^{f/f}$ [2] and *Crebbp*$^{f/f}$ [3] mice were crossed for the purpose of obtaining double or triple transgenics. Animals were housed according to the Spanish and European regulations and the experiments were approved by the Institutional Animal Care and Use Committee. Mice were caged in a 12-12 hour cycle (8:00-20:00) with the food and water available *ad libitum*. CaMKIIα-creERT2 carrying mouse lines were maintained in their standard housing without interruption for 3-4 months in order for them to fully develop the central nervous system. Then they were treated with tamoxifen (TMX) as described before [4], allowing for a complete elimination of either p300, CBP or both of these proteins. Mice heterozygous for the recombinase presented a complete elimination of a given protein (**Fig. 1b** and **Fig. S1a-b**) and their Cre$^-$ littermates were treated as controls. Animals in a very poor state or showing signs of pain were immediately sacrificed. To improve the well-being of the mice lacking both KAT3, we provided them with several high-protein food pellets (Teklad Global Diets® 2919, Envigo) always available on the bedding.

*Lentiviral production and plasmid constructs*

Lentiviral particles (LV) were produced according to the description in [5]. The following lentiviral constructs were acquired from Addgene: LV-CRE-GFP (Addgene #20781), LV-CRE-mCherry (Addgene #27546) and LV-RFP (Addgene #17619), used to label infected neurons and astrocytes; dCas9-KAT (Addgene #61357), LV-dCas9-KAT (Addgene #83889) and LV-U6gRNA-Cherry (Addgene #85708) used in epi-editing experiments; and LV-phND2-

N174 (Addgene #31822) used to overexpress human NeuroD2. All the designed gRNAs were cloned in the LV-U6gRNA-Cherry. The locations of gRNAs were selected based on the CBP, p300 and H3K27ac enrichment in the chromatin of dKAT3-iKOs mice. To produce LV-CREw/oGFP we digested LV-CRE-GFP with XbaI and BsrGI to remove GFP. As a control for Cre recombinase transduction, we infected with a synapsin promoter-bearing lentiviral vector derived from LenLox 3.7, LV-syn-syn-GFP [6]. The chimeric construct encoding CBP fragments were subcloned by PCR from full-length CBP: N-terminus-CBP-vhhGFP4 spans amino acids (aa) 1-1,098; GFP-C-terminus-CBP spans aa 1,099-2,441; NLS-KATGFP includes a sequence encoding a nuclear localization signal (KKKRKVD) fused to aa 1,088-1,758 of CBP and to full-length GFP. pDsRed-Express2-C1 (Clontech) was used in the analysis of neuronal morphology in hippocampal cultures.

*Primary cultures, lentiviral production and infection*

Primary hippocampal and cortical cultures were prepared as described in [7], with some modifications. *Crebbp*$^{f/f}$::*Ep300*$^{f/f}$ or *Crebbp*$^{f/f}$::*Ep300*$^{f/f}$::CAG/loxP/STOP/loxP /tdTomato embryos were dissected, dissociated, pooled together and plated in 24-well plates at $0.11x10^6$ neurons/well. After 24 hours, the primary cultures were infected with the corresponding virus (day *in vitro* 1, DIV1). In the experiments preventing CBP loss, neuronal cultures were co-infected at DIV1 with the indicated viruses and the cultures were processed after 11 days. In rescue experiments, neurons were first infected with a Cre recombinase expressing LV at DIV1, 11 days later the same neurons were transfected with the indicated vectors and the cultures were fixed at DIV13. To produce the astrocyte primary cultures, astrocytes were isolated from cortices of P1-P3 *Crebbp*$^{f/f}$*::Ep300*$^{f/f}$ pups. The cortices were dissected, cut into pieces and washed twice with HBBS 1x (Lonza BE10547F). The tissue was disrupted with a pasteur pipette with rounded edges in 2 ml of complete medium (DMED from Gibco, 21969-035 plus 10%FBS and 1% penicillin/streptomycin). The homogenate was filtered with a 70 µm cell strainer (Falcon #352350), centrifuged and resuspended in complete medium. The astrocytes from two pups were pooled together and plated in 24-well plates. Astrocytes cultures were infected at DIV10 with LV-CRE-mCherry or LV-RFP as a control, and fixed with PFA 4% at DIV17 or DIV30. In all immunohistochemistry experiments cells were cultured on glass coverslips coated with poly-lysine. N = 2 wells per PNC in 3 independent experiments.

*RNA isolation, RT-PCR and transfection*

Total RNA from hippocampal or cortical cultures was extracted with TRI reagent (Sigma-Aldrich) and reverse transcribed to cDNA using the RevertAid First-Strand cDNA Synthesis kit (Fermentas). RT-PCR was performed in an Applied Biosystems 7300 Real-Time PCR unit using Eva Green RT-PCR reagent mix. The transfections were done using Lipofectamine 2000 (Invitrogen) at DIV12 and the cultures were fixed at DIV13 for morphology analysis and rescue experiments.

*Antibodies*

The following primary antibodies have been used in this study: anti-CBP, Santa Cruz sc-583 (IHC, ChIP); anti-CBP, Santa Cruz sc-369 (ICC); anti-CBP, Santa Cruz sc-7300 (IHC, ICC); anti-p300, Santa Cruz sc-585 (IHC,

ICC, ChIP); anti-NeuroD2, Abcam ab109406 (ICC); anti-H2Aac, [8] (IHC); anti-H2Bac, [8] (IHC, ICC); anti-H3K9,14ac, [8] (IHC); anti-H3K27ac, Abcam ab4729 (IHC, ICC, ChIP); anti-H4ac, [8] (IHC); anti-NeuN, Millipore MAB377 (IHC, FANS); anti-Hpca, Abcam ab24560 (IHC, ICC); anti-CaMKIV, BD Transduction Laboratories C28420 (IHC); anti-Cleaved-Cas3, Cell Signalling #9661 (IHC); anti-Fos, Synaptic Systems #226004 (IHC); anti-mCherry/dsRed, Clontech 632496 (ICC); anti-GFP, Aves Labs GFP-1020 (IHC, ICC); anti-GFAP, Sigma G9269 (IHC, ICC); anti-H2A.Xγ, Abcam ab2893 (IHC); anti –GFP, Aves Labs GFP-1020 (IHC, ICC).

*Oligonucleotides*

The following oligonucleotides have been used in this study:

| **Target** | **Type** | **Forward** | **Reverse** |
|---|---|---|---|
| Gapdh | RT-qPCR primer pair for detection of mRNA | CTTCACCACCATGGA GAAGGC | CATGGACTGTGGTCA TGAGCC |
| CBP | RT-qPCR primer pair for detection of mRNA | TCAGCTCTTCCAACT TCCTTGG | AAGGAGGCGCTGCTG TAGGTAT |
| p300 | RT-qPCR primer pair for detection of mRNA | AAAAGACCGACGGAT GGAAAA | TTCTCGGCTAGGAGG TGATAGT |
| NeuN | RT-qPCR primer pair for detection of mRNA | GCAGTCGCGGTTGG AGTAGT | CGTTAAAAATGATCTC CACGTCTAAAAT |
| Gfap | RT-qPCR primer pair for detection of mRNA | GGACAACTTTGCACA GGACCTC | TCCAAATCCACACGA GCCA |
| Hpca | RT-qPCR primer pair for detection of mRNA | CTACATCAGCCGGGA GGAGAT | ATCTTGTAAATGGCCT GCACAA |
| Grik3 | RT-qPCR primer pair for detection of mRNA | GTACGGTGCTGTCAA GGACG | GGCCCACATCTTCTC AAAGGT |
| Gria1 | RT-qPCR primer pair for detection of mRNA | TGGAAGCAAGGACTC CGGAAGT | AACTCGATTAAGGCA ACCAGCATG |
| Kcnq2 | RT-qPCR primer pair for detection of mRNA | CGTTCATCTACCACG CCTACG | GCACAAGGCAGGAGA AGACTAAAA |
| Kcnq5 | RT-qPCR primer pair for detection of mRNA | TGGCTTCAAGTTGCC TCTTAATTC | CAAAGACAACGATCA TCACGAAC |
| mNeuroD2 | RT-qPCR primer pair for detection of mRNA | GAGATCCCTGAACCC ACGTT | TCATCTTGCGTTTCTT CGGC |
| hNeuroD2 | RT-qPCR primer pair for detection of mRNA | CTCGCCCGACCACG A | GCGCCGAGTAGTGCA TAGA |
| Fos Pr | ChIP-qPCR primer pair for Fos Promoter | CGCCCAGTGACGTA GGAAGT | GCAGTCGCGGTTGGA GTAGT |
| Bdnf Pr 3 | ChIP-qPCR primer pair | GACCAATCGAAGCTC | GGCACTGGGGTCAGA |

| | | | |
|---|---|---|---|
| | for Promoter 3 | AACCG | CATTA |
| Intergenic | ChIP-qPCR primer pair for control region ~75 kb upstream of *Bdnf* gene | CTACCGAGTGTTGAT TGCCGT | TGATGCAAGTGTCAA GCTCAATG |
| Control gRNA | Enhacer gRNA | GCTGGCCCTGATTTC GGGC | GGCCCGAAATCAGGG CCAG |
| Neurod2 Pr gRNA A | Promoter gRNA | GGGGTACCAGCCTC TATGCC | GGCATAGAGGCTGGT ACCCC |
| Neurod2 Pr gRNA B | Promoter gRNA | CCCCATTGTTCCCAT GTGGG | CCCACATGGGAACAA TGGGG |
| Neurod2 Pr gRNA C | Promoter gRNA | GAGATGCCACACTCG CTCCG | CGGAGCGAGTGTGG CATCTC |
| Neurod2 Pr gRNA D | Promoter gRNA | GTGGTGGGGGGGCG CTGCTT | AAGCAGCGCCCCCCC ACCAC |

*Behavioral testing*

All the behavioral testing has been performed using a mix of male and female mice. Animal's survival and well-being were monitored daily starting with the first day after TMX administration (Day 1) for at least one month (Day 30). The SHIRPA testing battery was performed as presented previously in [9]. *CamKIIa-CreERT2::Crebbp$^{f/f}$::Ep300$^{f/f}$*, *CamKIIa-CreERT2::Crebbp$^{f/f}$::Ep300$^{f/+}$* and *CamKIIa-CreERT2::Crebbp$^{f/+}$::Ep300$^{f/f}$* mice were first tested two days before TMX treatment and again three days after the last TMX injection (Day 12). SHIRPA phenotyping categories and detailed scoring can be found in the legend of **Table S1**.

*Stereotaxic surgeries and virus administration*

Mice where deeply anesthetized i.p. with a mixture of midazolam (5 mg/kg), medetomidine (1 mg/kg) and fentanyl (0.05 mg/kg) mixed in NaCl (0.9%) to a volume of 10-15 µl/g of body weight. As soon as the total loss of reflexes was observed, the animals were positioned in a digital stereotaxic frame (Stoelting). At this point the body temperature was constantly monitored and maintained during the surgery at 37ºC using electric blanket. Local anesthetic

(EMLA 25%, lidocaine/prilocaine, AstraZeneca) was applied on the ear bars and the ophthalmologic gel (Viscotears, Bausch + Lomb) was administered on the eyes to avoid the formation of ulcers. Once the cranium has been exposed, a hole was drilled in the calvaria in the location corresponding to the hippocampus. A glass capillary (Word Precision Instruments) containing the adeno-associated virus (AAV) was placed slowly in the coordinates of the hippocampal hilus (-2.0, +/-1.35, -1.95; in mm relative to bregma) and left in place for five minutes. 500 nl of either AAVs rAAV5-hSyn-GFP-Cre or rAAV5-hSyn-mCherry-Cre was injected (Vector Core at the University of North Carolina at Chapel Hill). The capillary was then left in place for another five minutes and withdrawn. After the surgery, anesthesia was reversed with subcutaneous atipamezole (0.02 mg/kg). Buprenorphine in food pellets (1 mg/ml) was placed in the home cages to reduce post-surgery pain and mice where monitored daily until fully recovered.

*In vivo electrophysiology*

Mice were deeply anaesthetized with 4% isoflurane (Isoflo®, Esteve Veterinaria S.A.) in 0.8 L/min oxygen and fixed in a stereotaxic setup (NARISHIGE Group) over a heating pad at 37°C. Isoflurane was kept at 1-2%, 0.8 L/min oxygen to maintain the anaesthesia. After checking the lack of reflexes, mice were placed and fixed in a stereotaxic frame (NARISHIGE Group). The skin of the head was cut, and the scalp and periosteum were separated. Two 1.8 mm Ø holes were made in the skull using a milling cutter (FST 18004-18, Fine Science Tools) attached to a cordless micro drill (58610V, Stoelting Co.) in the appropriated coordinates to introduce the electrodes. Then one bipolar stimulating electrode (10-15 kΩ, 325 μm Ø,

TM53CCNON, World Precision Instruments) was introduced in the perforant pathway (from bregma, in mm: -4.3 AP, +2.5 ML, +1.4 DV, 12$^{o}$ angle), and one recording probe (single shank, 50 µm contact spacing, 32 channels; NeuroNexus Technologies) was targeted to the hippocampus CA1 and dentate gyrus regions (from bregma, in mm: -2 AP, +1.5 ML, -2 DV). Recording and stimulating electrodes were implanted following stereotaxic standard procedures and optimized based on the online recording to have the best quality of the signal in the dentate gyrus and CA1, especially taking into account the typical evoked potential in dentate gyrus [10]. A custom-made Ag/AgCl wire was placed in contact with the skin and used as a ground. After optimizing the final position, the tissue was allowed to rest for 30 minutes before acquiring electrophysiological data. The position of the electrodes was confirmed *post mortem*. The stimulating electrode was connected to a pulse generator and current source (STG2004, Multichannel Systems) controlled by MC_Stimulus software (Multichannel Systems). Electrophysiological data from the recording probes were filtered (0.1-3 kHz), amplified and digitalized (20 kHz sampling rate for evoked potentials and 32 kHz for spontaneous activity recordings) and analysed off-line using the Spike2 software (Cambridge Electronic Design Limited) or MATLAB (MathWorks) using ICAofLFPs package [11, 12]. Stimulating and recording protocols entailed spontaneous recordings (5 minutes) and evoked potentials, that consisted of a classical Input-Output (IO) stimulation protocol (stimulation intensities of 0.05, 0.1, 0.2, 0.4, 0.6, 0.8, 1 and 1.2 mA). For evaluating the Excitatory Post-Synaptic Potential (EPSP) the deepest slope of the evoked potential in molecular layer (dentate gyrus) was measured. To reflect the Population Spike (PS) the amplitude of the spike recorded in hilus was measured. Data were averaged by animal, per stimulation intensity, and then by group***.*** Spontaneous activity signals coming from representative channels in dentate gyrus were selected to analyse the power of the frequency bands and the wavelet spectrum. Briefly, after down-sampling of spontaneous recordings to 2.5 KHz, the signals were filtered (high pass at 0.5 Hz and notch at 50 and 100) and then analysed to extract: a) its power density by frequency bands; b) the wavelet spectrum, using the Fourier Transformation or the Wavelet spectrum analysis, respectively, implemented in the MATLAB package ICAofLFPs. Single Unit Activity (SUA), as local activity reflex, was analysed using a supervised tool integrated in Spike2 software. Briefly, after applying a band pass (0.3-3 KHz, Butterworth digital filter), the different waveforms were extracted with intensity threshold set at ± $3^{-4}$ mV (to avoid noise) for all the putative SUA-spikes recorded nearby the recording electrode in the dentate gyrus or in CA1. After the complete scan of the electrophysiological signal, per area, we manually chose only those waveforms that clearly fit with the typical one reflecting neuronal activity (supervised procedure). The number of spikes was averaged by area, animal, and then by group to avoid an overestimation of the total *n* for the statistical comparisons.

*Histology and image processing*

Experimental and control animals were anesthetized using a mix of xylazine (Xilagesic, CALIER) and ketamine (Imalgene, MERIAL LABORATORIOS) and perfused transcardially, first with PBS (pH 7.4) to removed whole blood, then with a solution containing 4% paraformaldyhyde in PBS. Brains were perfused and postfixed overnight (4% paraformaldehyde) and subsequently cut on vibratome into 50 µm sections. Sections were used for immunohistochemistry (fluorescent and diaminobenzidine) or Nissl staining as previously described in [4]. Some antibodies used for the fluorescent immunohistochemistry required a 30 min antigen retrieval in 80°C sodium citrate buffer (10mM Sodium Citrate, 0.05% Tween 20, pH 6.0). The Golgi-Cox impregnation has been performed using the FD Rapid GolgiStain™ Kit (FD NeuroTechnologies, Inc.). To this end, animals were anesthetized using a mix of xylazine and ketamine to prevent any head damage and sacrificed through cervical dislocation. The brains were instantly removed from the skull, rinsed very briefly with double distilled water in order to remove the excess of blood and placed in the impregnation solution in the room temperature. The solution was changed to a fresh one after the first 24 hours. After 10 days the brains were immersed in solution C for 72 hours, after which it was changed to a fresh solution C for further 24 hours. Subsequently, the brains were carefully cut on vibratome into 100 µm sections in a 1:1 mix of PBS and solution C. These sections were next mounted on gelatin-coated slides, revealed using solutions D and E, dehydrated using ethanol and xylene and covered using Neo-Mount® (Merck). This protocol resulted in a sparsely marked neurons along the entire section of the brain, ideal for reconstructing the neuronal morphology (see next section). Samples marked using Golgi technique were visualized under a bright field microscope with a motorized stage. Dentate gyrus granule neurons with mostly intact dendritic trees were traced and subsequently reconstructed using the Neurolucida software (MBF Bioscience). The reconstructions were used to perform a Sholl analysis, where the number of intersections was calculated every 2 µm starting from 5 µm from the soma. The thickness of

CA1 sub-regions was quantified based on the Nissl staining (sagittal cut) bright field images made with a 2.5x objective. In Fiji software four lines were drawn perpendicularly to the CA1 and the thickness of *stratum pyramidale* or *stratum radiatum* was measured along each of the lines. The average value was the final measurement for each animal. For electron microscopy (EM) experiments, mice were anesthetized and perfused as described for the immunohistochemistry with the addition of 2,5% glutaraldehyde in the fixation solution. Then brains were then cut on vibratome to 100 µm slices. Selected slices with dorsal hippocampus were postfixed with 1% osmium tetroxide for 1h at room temperature. Dehydration was performed by incubating the slices in increasing ethanol concentrations and in pure propylene oxide. During dehydration, tissue was stained with 1% uranyl acetate in 70% ethanol. Slices were then embedded in the Epon resin between two Aclar sheets. After polymerization, Cornu Ammonis (CA1), *stratum radiatum* and dentate gyrus regions fragments were cut out and stuck to an empty block of resin. Next, 75 nm sections were prepared and post stained with uranyl acetate and Reynold's lead citrate. Electron micrographs were taken with JEM 1400 transmission electron microscope at 80 kV (JEOL Ltd. 2008). Quantification of synapses and heterochromatin clumps number were performed using Fiji software with the *Cell Counter* plugin or macro based on the *Analyze Particles* function respectively.

*Fluorescence-activated nuclear sorting (FANS)*

Experimental mice were sacrificed by cervical dislocation and the hippocampal tissue was extracted from the brains. This tissue was subsequently homogenized using a douncer tissue grinder (Kontes® 2ml) in a buffer containing 0.5% IGEPAL and filtered on a 35μm nylon mesh (Falcon #352235). The resulting suspension of hippocampal nuclei was stained using mouse anti-NeuN antibody, anti-mouse Alexa 647 antibody and DAPI. At this point three independent samples were pooled to form a single replicate which was centrifuged in Optiprep (MERCK) gradient. Nuclei purified this way were sorted using BD FACS Aria III Flow Cytometer by their size (FSC), complexity (SSC), DAPI and NeuN-Alexa647. dKAT3-ifKO samples contained two populations of neuronal nuclei: (I) strongly immunofluorescent for NeuN, which did not undergo Cre recombination; and (II) weakly immunofluorescent for NeuN, which still expressed higher NeuN levels than non-neuronal cell-types. Specific gate was set up in the flow cytometer to isolate only the second group of dKAT3-ifKO nuclei for the ATAC-seq experiments (NeuN+, **Fig. 4c**). Approximately 50,000 nuclei were obtained in each case (41.5% of events in control and 46.45% in dKAT3-ifKO).

*Genomic data processing and access*

All sequenced datasets adapters were trimmed using cutadapt v1.18 [13] and aligned to mm10. Only reads longer than 25 bp, with mapq > 30 and mapping to nuclear chromosomes were used for the posterior analysis. Data was processed with the extensive use of custom R scripts (R version 3.5.1, 2018), Samtools v1.9 [14], bedtools v2.27.1 [15] and DeepTools v3.2.0 [16]. Whole genome alignments were normalized to 10xRPM (read per 10 million sequenced reads) and visualized using IGV v2.5.0 [17].

*mRNA-seq:* Extraction of RNA from the hippocampal tissue was performed using TRI-reagent (MERCK) as previously described [18]. Resulting total RNA was treated with DNase I (Qiagen) and its quality was confirmed using nanodrop, Bioanalyzer and RT-PCR. Three independent samples were prepared for both Control and dKAT3-ifKO mice. Each sample, which corresponded to a single mouse, was used to prepare a polyA library and sequenced on the HiSeq 2500 sequencer (Illumina, Inc). Reads were aligned with HISAT2 v2.1.0 [19] to mm10 mouse genome. Mapped reads were annotated to genes from Ensemble (GRCm38.89) and quantified using HTseq v0.11.1 [20]. Differential expression analysis was performed using the Bioconductor package DESeq2 v1.10.0 [21]. Genes with FDR < 0.05 and log2FC > +/- 1 were considered significantly deregulated. GO terms where analyzed using the Bioconductor package *Gostats*. Tissue specific gene expression was taken from GTEx [22], selecting *Brain-Hippocampus*, *Heart-Left Ventricle*, *Liver* and *Lung*.

*ATAC-seq:* The Assay for the Transposase Accessible Chromatin followed by high-throughput sequencing (*ATAC-seq*) was performed as described in [23, 24]. Briefly, sorted neuronal nuclei were centrifuged and resuspended in the transposase reaction mix (TD buffer and Tn5 transposase, Illumina). The mix was placed in 37°C for 30 min and immediately after the DNA was extracted using Qiagen MinElute PCR Purification Kit. A DNA library was then prepared using Custom Nextera PCR primers 1 and 2. We monitored the saturation of the library using an RT-PCR and afterwards extracted the DNA using the kit. This final DNA extract was sequenced using HiSeq 2500 sequencer (Illumina, Inc). Paired-end reads were aligned with Bowtie2 v2.3.4.2 [25] to mm10 mouse genome. Duplicated reads were removed with Picardtools v2.18.21 (https://broadinstitute.github.io/picard/). Only paired reads were used for posterior analysis. Peak calling was performed with MACS2 v2.1.1 [26]. Following ENCODE recommendations, to filter the most reproducible and better quality peaks, called peaks from ChIP replicates were subjected to Irreproducible Discovery Rate (IDR) selection and only peaks with IDR < 0.15 were taken for downstream analysis. DARs analysis was performed using the Bioconductor package DiffBind [27]. Regions with FDR < 0.05 and log2FC > +/-1 were considered significantly regulated. DARs were annotated to closest genes from Ensembl (GRCm38.89) using the Bioconductor package ChIPpeakAnno [28]. Predictive relationship of DARs to gene expression changes was performed using BETA v1.0.7 [29] using as reference genes from Ensembl (GRCm38.89). Motif analysis of ATAC regions was performed using MEME-suite [30]. For digital footprint we chose the ATAC dedicated software HINT [31]. The comparison of lost ATAC-seq signal and Eigen vector values from the three differentiation stages explored in [32] (GSE96107) and our recent study in mature excitatory neurons [24] (GSE125068) was done using FPKM values from each dataset.

*ChIP-assay and ChIP-seq*

H3K27ac chromatin immunoprecipitation was performed as previously described [33]. The CBP and p300 ChIP experiments required the following adjustments: Minced hippocampal tissue was fixed in 1% PFA for 30 min in 37°C (as suggested in [34]), which allowed for crosslinking of the KAT3 cofactors to the DNA-binding proteins and to the DNA itself. As the KAT3 proteins disintegrate almost completely after a few rounds of sonication in 1% SDS (unpublished data of Barco lab), the sonication buffer was changed to one containing 0.1% SDS, 1% IGEPAL (Sigma-Aldrich) and 0.5% sodium deoxycholate. Additional 10 sonication cycles of 30''on/30''off were added to fragment the highly fixed DNA sufficiently for sequencing. Reads were aligned with Bowtie2 v2.3.4.2 [25] to mm10 mouse genome. Peak calling was performed with MACS2 v2.1.1 [26]. Following ENCODE recommendations, we used IDR < 0.05 to avoid false positives and retrieve the most reliable peaks. Heatmaps were performed with *DeepTools*. Circos plot was drawn with the R package *Circlice*. Differential protein binding analysis between ChIP controls and dKAT3-ifKO was performed using the Bioconductor package DiffBind [27]. Regions with FDR < 0.05 and log2FC > ±1 were considered significantly deregulated. Annotation and handling of ChIP peaks was performed with Bioconductor packages ChIPseeker [35] and ChIPpeakAnno [28]. ChIP peaks were annotated to closest genes from Ensembl (GRCm38.89). Motif analysis of regions occupied by ChIP peaks was performed using MEME-suite [30]. For the classification of regulatory regions categorized by genomic features, neuronal and pancellular KAT3 peaks were split according to H3K27ac enrichment and H3K4me1/H3K4me3 content (information of H3K4me1 ChIP-seq from ENCODE (ENCFF545CTN) and H3K4me3 ChIP-seq from [18]). Most H3K4me3-rich regions lie up to 1 kb from the TSSs corresponding to what could consider promoters of active genes, while H3K4me1-rich peaks preferentially locate into introns and intergenic regions and were labeled as enhancers. We defined as neuronal enhancers those regions that contain neuronal peaks for KAT3 binding, ATAC-seq, H3K27ac, enriched in H3K4me1 and located in introns or intergenic regions. Peaks that contain KAT3, ATAC-seq, H3K27ac and H3K4me3 enrichment located at promoters were labelled as active promoters. To retrieve putative neuronal super-enhancers we used a similar criteria than Whyte *et al.,* [36], H3K27ac regions closer than 5 Kb were stitched together, then the previously described neuronal enhancers were tested. Regions containing neuronal enhancers were carried over for further analyses. From those, we retrieved the regions longer than 5 kb (the length at which we see that gene expression deviates from linear correlation) and associated with genes with a basal gene expression one order of magnitude higher than average (100 RPKM). The regions that satisfied both conditions were labeled as neuronal super-enhancers (SEnh). The information of Neurod2 ChIP-seq is extracted from [37]**.** The Phenotype analysis was performed using the WEB-based application GEne SeT AnaLysis Toolkit (http://www.webgestalt.org/). Murine (C57BL/6) p300 ChIP-seq at P0 data was taken from ENCODE for the tissues heart (ENCSR777VNA), liver (ENCSR765RPR), and lung (ENCSR527DME).

*Single-nucleus RNA sequencing and analysis*

For the single-nucleus RNA-seq experiment, dKAT3-ifKO mice were sacrificed either 2 weeks (2w) or 1 month (1m) after the TMX administration. The two hippocampi of each mouse were dissected in cold PBS and transferred to a dounce homogeneizer containing 1 ml of ice cold MACS buffer (0.5% BSA, 2 mM EDTA, PBS 1x) and were homogenized 15-12 times with the pestle. The cell suspension was transferred to a 2 ml tube and centrifuge 15 min at 500g and 4ºC. Cell pellets were resuspended in 2 ml of lysis buffer (10 mM Tris-HCl, 10 mM NaCl, 3 mM $MgCl_2$, 0,1% IGEPAL) and kept 5 min on ice. Samples were then spun down at 500g for 30 min in a pre-chilled centrifuge. The nuclei pellet was resuspended in PBS 1x 1% BSA and sorted on a BD FACS Aria III. 15,000 nuclei per sample (pool of 2 animals) were loaded into the single cell A Chip and then generation of barcode-containing partitions were carried out with the Chromium Controller (10X Genomics). Chromium Single Cell 3' Library & Gel Bead Kit v2 was employed for post-GEM-RT clean-up, cDNA amplification and the generation of barcoded (ChromiumTM i7 Multiplex Kit) libraries. Libraries were sequenced to an average depth of 290-310 Million reads per sample on an Illumina HiSeq2500 sequencer according to manufacturer instructions. Quality control of sequenced reads was performed using FastQC (Babraham Institute). Sequenced samples were processed using the Cell Ranger v2.2.0 pipeline (10x Genomics) and aligned to the CRGm38 (mm10) mouse reference genome customized to count reads in introns (pre-mRNA) (gene annotation version 94). We retrieved 1,791 (control), 1,133 (dKAT3-ifKO 2w), and 1,465 (dKAT3-ifKO 1m) high quality nuclei per sample. Mean reads per nucleus were 172,879 (control), 271,535 (dKAT3-ifKO 2w), and 197,840 (dKAT3-ifKO 1m). Single-nucleus RNA-seq data were subsequently pre-processed and further analysed in R using Seurat v2.3.4 [38, 39]. Filtering parameters were as follows: genes, nCell <5; cells, nGene <200. Data were then normalized using global-scaling normalization (method: LogNormalize, scale.factor=10.000). To identify major cell populations in the dorsal hippocampus of adult mice, control and dKAT3-ifKO datasets were analysed separately. Highly variable genes (HVGs) were detected using *FindVariableGenes* function with default parameters. Then, normalized counts on HVGs were scaled and centred using *ScaleData* function with default parameters. Principal component analysis (PCA) was performed over the first ranked 1,000 HVGs, and cluster detection was carried out with Louvain algorithm using 20 first PCA dimensions at resolution = 0.6 (the default and the optimal according to cell number, data dispersion and co-expression of previously reported cell markers). Visualization and embedding were performed using tSNE and UMAP over PCA using the 20 first PCA dimensions. UMAP plots of gene expression show normalized count (UMIs) per nucleus. The equalized expression between fixed percentiles was plotted according to the following criteria: the minimum expression was adjusted to 25% and the maximum expression was adjusted to 95% in all expression plots. For longitudinal analysis, datasets from the 3 conditions were merged and HVGs were identified for each dataset as above indicated. Only HVGs that were detected in all datasets were used to perform visualization and embedding as described above. Clustering was performed on merged dataset from 3 conditions and populations were identified combining these results with clustering information obtained in control and dKAT3-ifKO 1m datasets separately, together with co-expression of population markers. Differential expression analysis (DEA) was used to identify population gene markers. For DEA, the nuclei of each population were contrasted against all the other nuclei in the merged dataset using Wilcoxon Rank Sum test on normalized counts.

*Statistical analysis*

All the statistics in the following work have been done using RStudio, GraphPad Prism or MATLAB, depending on the experiment. Statistical tests used in the study are indicated either in the Figure or in the corresponding legend. All statistical tests used in this study were two-sided, except for the Hypergeometric test. For a comparison of two groups, each group was first tested for the normality using Shapiro-Wilks test. If normality assumption was not violated a t-test was performed. If the normality null hypothesis was rejected, Mann-Whitney U test was performed. In the analysis of the SHIRPA paradigm, Fisher Exact Test was used for the categories carrying just two possible outcomes (indicated in **Table S1**). For multiple testing of the same sample, p-values were corrected using Bonferroni method. For two factor comparisons, Two-Way ANOVA was used. In all bar plots the height represents the mean and the error bars the standard error of mean (SEM). *ns:* non-significant, *: p-value < 0.05, **: p-value < 0.01; ***: p-value < 0.001.

*Data access*

Genomic data sets can be accessed at the GEO public repository (GSE133018).

#### Supplementary figures legends

**Figure S1 related to Figure 1. Conditional ablation of KAT3 genes in forebrain excitatory neurons. a.** DAB immunohistochemistry images showing CBP/p300 loss in forebrain neurons of a corresponding knockout. Notice no change in the expression of the protein in the cerebella and a complete elimination of the protein in the ifKO hippocampi (higher magnifications on the right). **b.** Immunohistochemistry against CBP and p300 in the CA1 region of wild type, CBP-ifKO and p300-ifKO mice. Dashed line contour indicates the position of the nuclei based on the DAPI signal. A full ablation of both proteins can be observed in pyramidal and granule neurons (not shown) in the hippocampus. Scale: 10 µm. **c.** Score of dKAT3-ifKO and control littermates in several significantly affected SHIRPA categories. The animals were scored twice: right before starting the TMX treatment ("0") and 12 days after the first TMX administration ("12"; Control, n = 6; dKAT3-ifKO, n = 8). Each category is scored according to a specific scale. The details regarding the scoring rules can be found in the Methods and **Table S1**. **d.** Locomotor activity in the open field as measured by the number of entered squares. Values for control mice are shown in black, CaMKIIα-CreERT2::Crebbp$^{f/f}$::Ep300$^{f/f}$ in red, *CaMKIIα-CreERT2::Crebbp*$^{f/+}$*::Ep300*$^{f/f}$ in magenta, and *CaMKIIα-CreERT2::Crebbp*$^{f/f}$*::Ep300*$^{f/+}$ in orange. The left panels show the results before TMX treatment and the right panels,12 days after the first TMX administration. **e.** Survival curve during the first month after TMX for *CaMKIIα-CreERT2::Crebbp*$^{f/+}$*::Ep300*$^{f/f}$ *and CaMKIIα-CreERT2::Crebbp*$^{f/f}$*::Ep300*$^{f/+}$ mice.

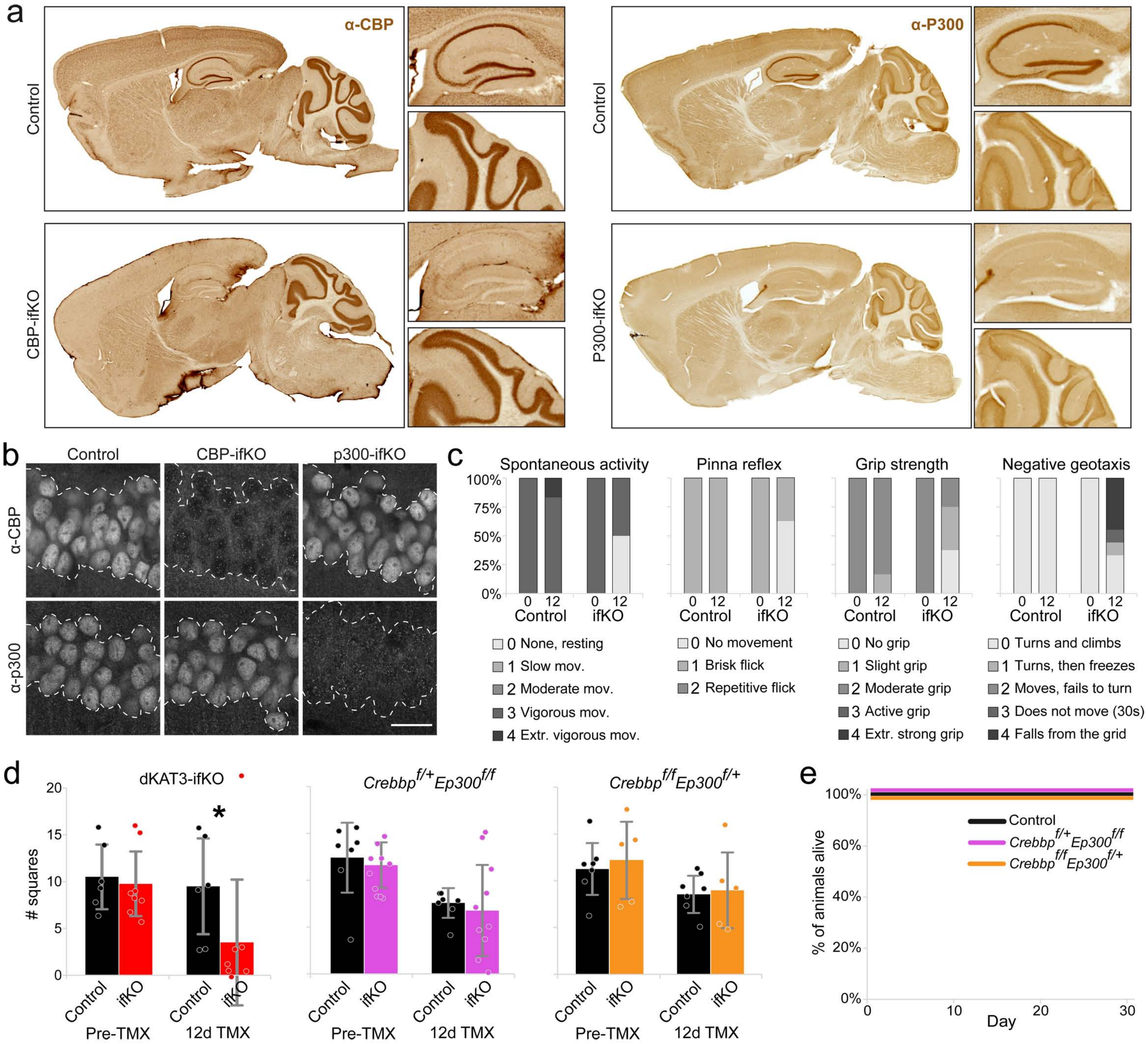

a
α-CBP
Control
CBP-ifKO
α-P300
Control
P300-ifKO
b
Control
CBP-ifKO
p300-ifKO
α-CBP
α-p300
c
Spontaneous activity
Pinna reflex
Grip strength
Negative geotaxis
100%
75%
50%
25%
0%
0 12
Control
ifKO
0 None, resting
1 Slow mov.
2 Moderate mov.
3 Vigorous mov.
4 Extr. vigorous mov.
0 No movement
1 Brisk flick
2 Repetitive flick
0 No grip
1 Slight grip
2 Moderate grip
3 Active grip
4 Extr. strong grip
0 Turns and climbs
1 Turns, then freezes
2 Moves, fails to turn
3 Does not move (30s)
4 Falls from the grid
d
dKAT3-ifKO
Crebbp^f/+ Ep300^f/f
Crebbp^f/f Ep300^f/+
### squares
20
15
10
5
0
*
Control
ifKO
Pre-TMX
12d TMX
e
100%
80%
60%
40%
20%
0%
% of animals alive
Control
Crebbp^f/+ Ep300^f/f
Crebbp^f/f Ep300^f/+
0
10
20
30
Day

**Figure S2 related to Figure 1. Dramatic reduction in the number of synapses and spontaneous and evoked neuronal activity in dKAT3-ifKOs. a.** Nissl staining showing normal gross anatomy and histology 1 month after single and combined CBP/p300 ablation. **b.** Immunostaining against GFAP shows no gliosis in the hippocampus of CBP-ifKOs, p300-ifKOs and in most dKAT3-ifKOs. Although we observed moderate active gliosis in a subset of dKAT3-ifKOs (less than 10%), this was not a general feature of dKAT3-ifKO brains. Instead, gliosis may rather result from the occasional episodes of epilepsy observed in some mice in the first days after gene ablation. Scale: 50 µm. **c.** Wavelet spectra of dentate gyrus Local Field Potentials (LFPs) from control (left) and mutant (right) mice. LFP signal from the same channel is presented as a white line in the upper part of each panel. A.U. – arbitrary unit. **d.** Dentate gyrus LFPs band-power in the delta (0-4 Hz), theta (4–8 Hz), alpha (8–13 Hz), beta (13–30 Hz), and gamma (30–125 Hz) frequency bands. **e.** Left: Quantification of EPSP slope of the stimulus-response curve recorded in dentate gyrus after perforant pathway stimulation for control (black) and mutant (red) mice. Right: representative EPSPs evoked potentials. Note the six-time larger stimulation intensity applied to dKAT3-ifKOs.

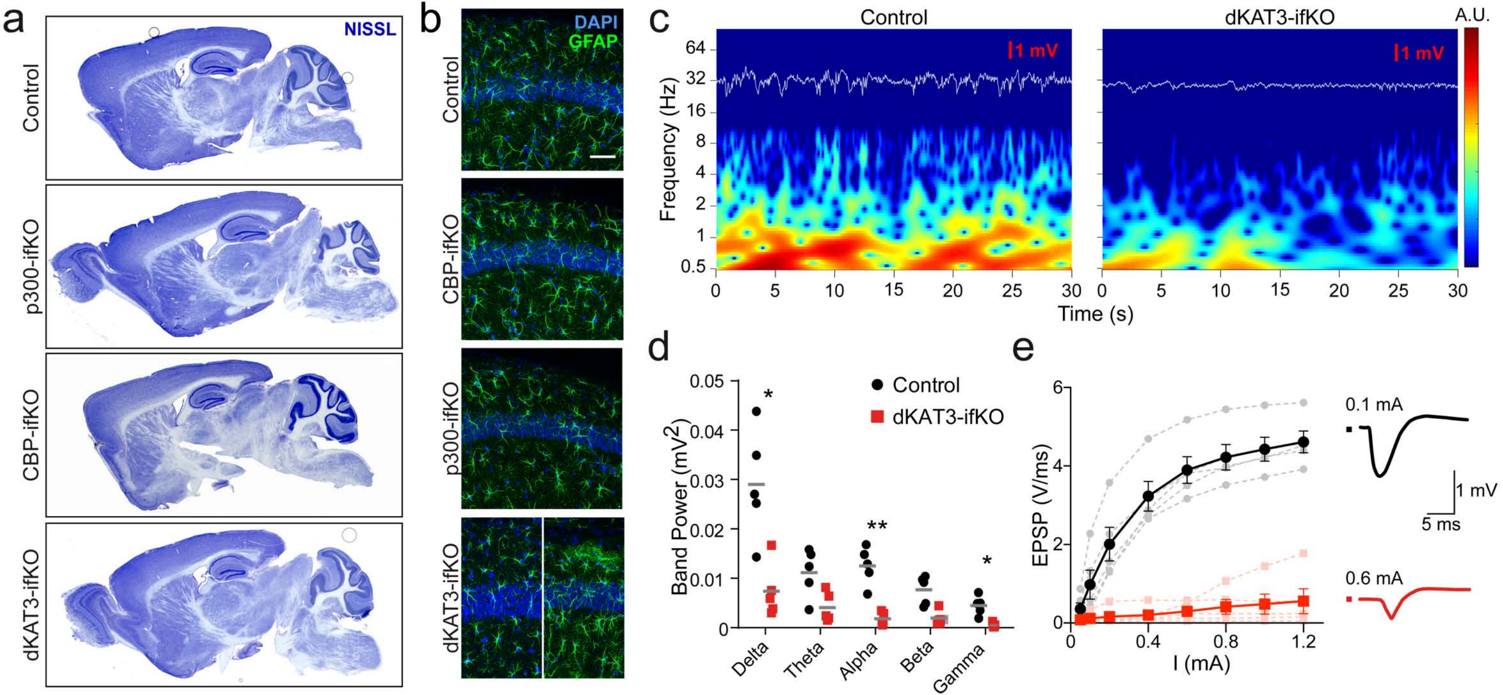
a
NISSL
Control
p300-ifKO
CBP-ifKO
dKAT3-ifKO
b
DAPI
GFAP
Control
CBP-ifKO
p300-ifKO
dKAT3-ifKO
c
Control
dKAT3-ifKO
A.U.
1 mV
1 mV
Frequency (Hz)
64
32
16
8
4
2
1
0.5
0
5
10
15
20
25
30
Time (s)
d
Control
dKAT3-ifKO
Band Power (mV2)
0.05
0.04
0.03
0.02
0.01
0
*
**
*
Delta
Theta
Alpha
Beta
Gamma
e
EPSP (V/ms)
6
4
2
0
0
0.4
0.8
1.2
I (mA)
0.1 mA
0.6 mA
1 mV
5 ms

**Figure S3 related to Figure 1. The double ablation of CBP and p300 does not cause neuronal loss. a**. TUNEL staining images (top) and quantification (below) show no cell death either in the brain of mice from either genotype. A section treated with DNase I was used as a positive control. Scale: 200 µm. **b.** Immunostaining against phosphorylated (pS139) histone H2A.X gamma, a marker of DNA damage related to cell death. Scale: 30 µm. **c.** Electron microscopy (EM) images of CA1 nuclei in control and dKAT3-ifKO mice 1 month after TMX. Scale: 2 µm. **d.** Quantification of nuclear features in EM images 1 and 2 months after TMX. Left: number of large (0.1 - 1 µm, top) and small (0.02 - 0.1 µm, bottom) electrodense granules per nucleus. Right: number of nuclear envelope invaginations per nucleus. **e**. The upper scheme describes the AAV-Cre-GFP stereotaxic injection experiment. AAV-Cre-GFP virus was injected unilaterally into the DG of adult *Crebbp*$^{f/f}$*::Ep300*$^{f/f}$ mice. **f.** Immunostainings showing the loss of both CBP and p300 in AAV-cre transduced Crebbp$^{f/f}$::Ep300$^{f/f}$ mice. Scale: 200 µm. **g.** Immunohistochemistry for Cleaved Cas3 in AAV-cre-transduced mice 1 and 2 months after infection. Scale: 200 µm.

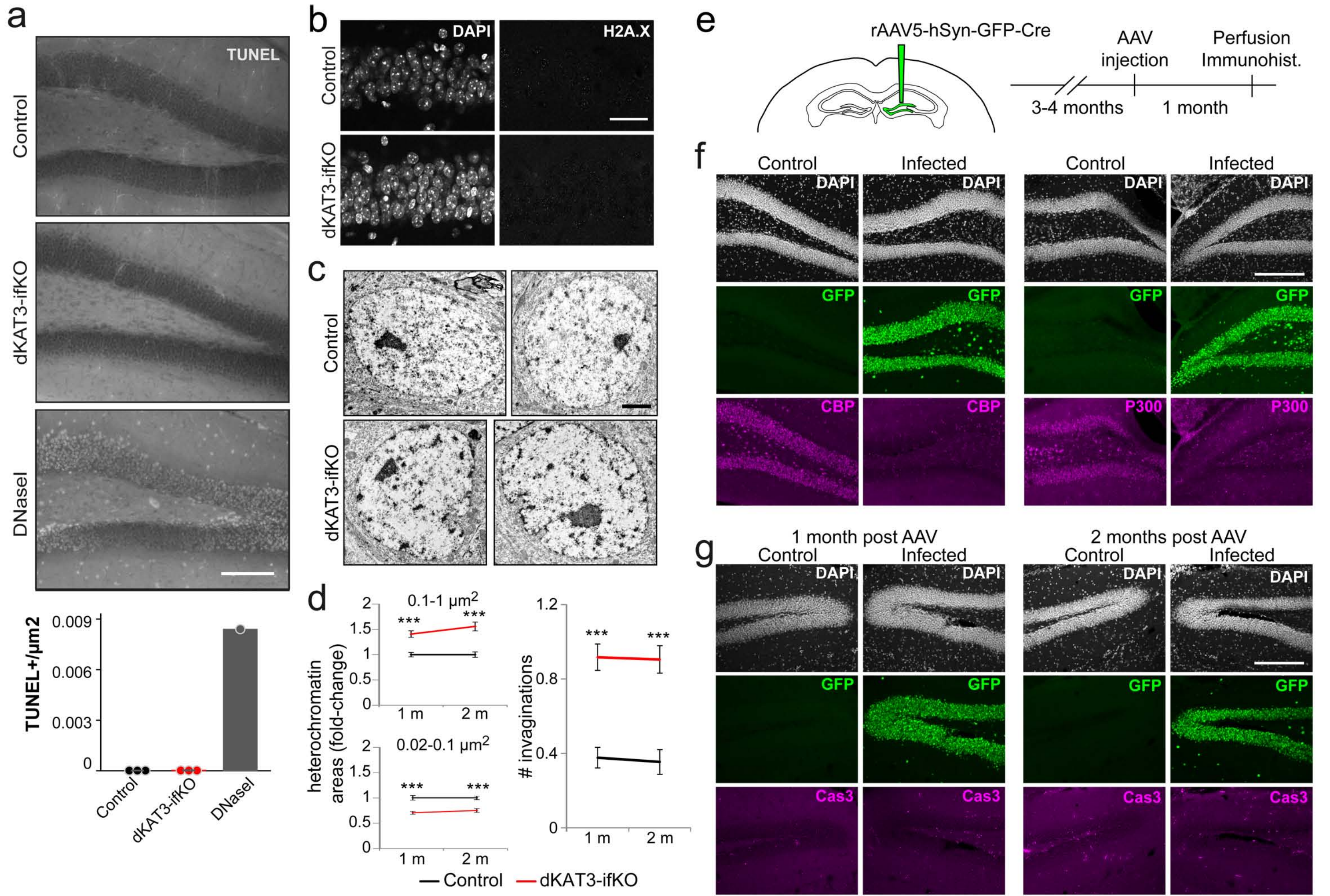
a
TUNEL
Control
dKAT3-ifKO
DNaseI
TUNEL+/µm2
0.009
0.006
0.003
0
Control
dKAT3-ifKO
DNaseI
b
DAPI
H2A.X
Control
dKAT3-ifKO
c
Control
dKAT3-ifKO
d
0.1-1 µm2
0.02-0.1 µm2
heterochromatin areas (fold-change)
***
1 m
2 m
### invaginations
1.2
0.8
0.4
0
Control
dKAT3-ifKO
e
rAAV5-hSyn-GFP-Cre
AAV injection
Perfusion Immunohist.
3-4 months
1 month
f
Control
Infected
DAPI
GFP
CBP
P300
g
1 month post AAV
2 months post AAV
Control
Infected
DAPI
GFP
Cas3

**Figure S4 related to Figure 2. Hippocampal cells lacking KAT3 fail to express neuronal-specific genes. a.** Principal component analysis plot of the RNA-seq experiment comparing hippocampal RNA of dKAT3-ifKO and control littermates. **b.** The genesets related with apoptosis and cell death (according to Gene Ontology) are not differentially expressed in dKAT3-ifKOs. Each dot represents a single gene. Red: upregulated genes; blue: downregulated genes; grey: no change. The change of the most severely downregulated positive regulator of neuronal death, *Harakiri* (*Hrk*), is labeled. **c.** RNA-seq profile for *Hrk* in dKAT3-ifKOs and control littermates. **d.** Examples of RNA-seq profiles for two housekeeping genes: *Ppia* (peptidylprolyl isomerase A) and *Pgk1* (phosphoglycerate kinase 1). **e.** Immunostaining against CBP and NeuN focusing in a rare field of the dentate gyrus containing cells in which there was no recombination and NeuN expression was maintained (arrowheads). Scale: 50 µm. **f.** RT-PCR gene expression analysis in the hippocampus of CaMKIIα-CreERT2::*Crebbp*$^{f/+}$::*Ep300*$^{f/f}$ and CaMKIIα-CreERT2::*Crebbp*$^{f/f}$::*Ep300*$^{f/+}$ mice. The expression of target genes was decreased more than 2-fold in the dKAT3-ifKO hippocampi (**Table S2**). Notice that only *Kcnq5* shows a slight downregulation in CaMKIIα-CreERT2::*Crebbp*$^{f/f}$::*Ep300*$^{f/+}$ tissue.

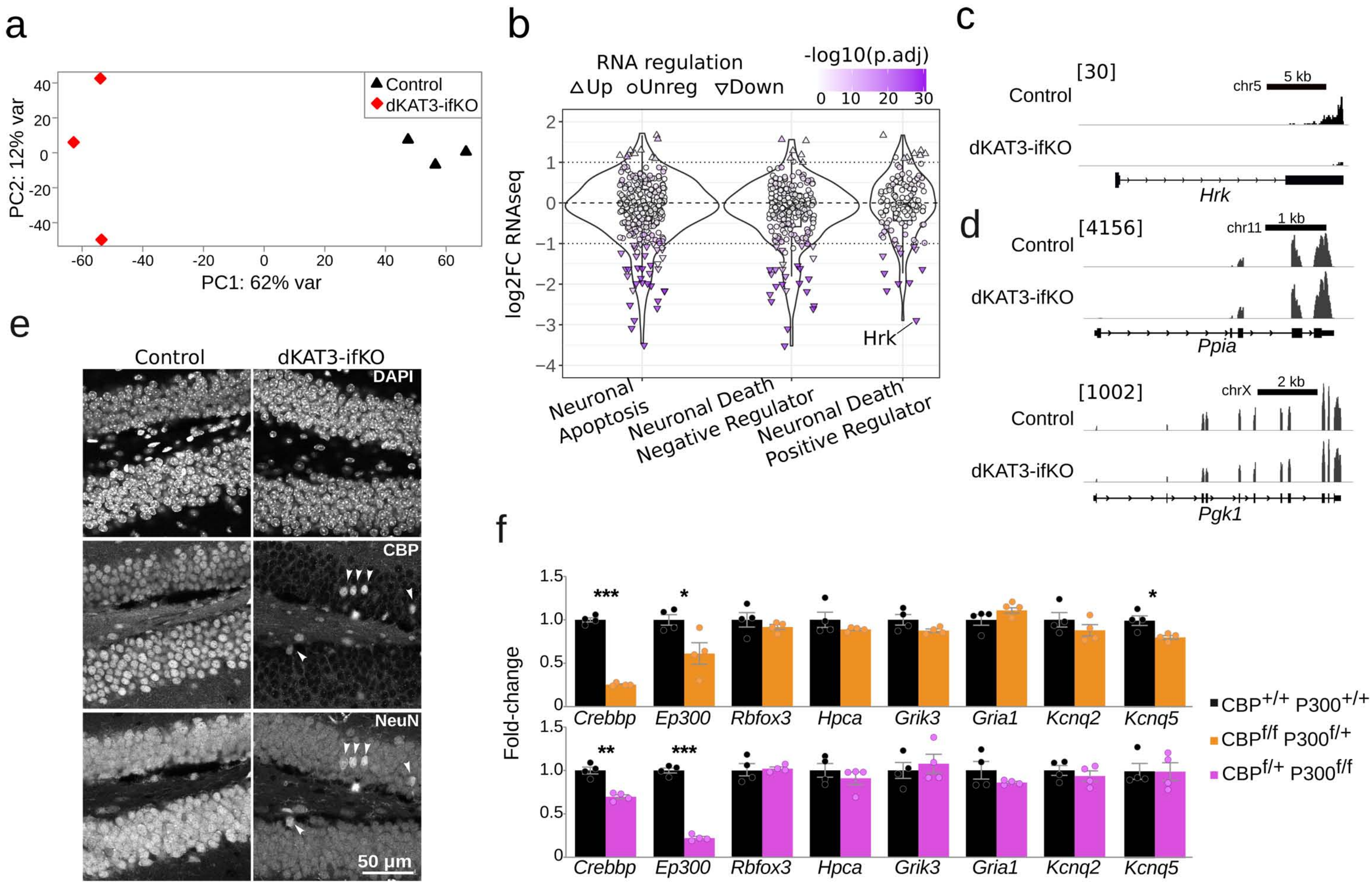
a
PC2: 12% var
PC1: 62% var
Control
dKAT3-ifKO
b
RNA regulation
Up
Unreg
Down
-log10(p.adj)
log2FC RNAseq
Hrk
Neuronal Apoptosis
Neuronal Death Negative Regulator
Neuronal Death Positive Regulator
c
[30]
Control
dKAT3-ifKO
chr5
5 kb
Hrk
d
[4156]
Control
dKAT3-ifKO
chr11
1 kb
Ppia
[1002]
Control
dKAT3-ifKO
chrX
2 kb
Pgk1
e
Control
dKAT3-ifKO
DAPI
CBP
NeuN
50 μm
f
Fold-change
Crebbp
Ep300
Rbfox3
Hpca
Grik3
Gria1
Kcnq2
Kcnq5
CBP+/+ P300+/+
CBPf/f P300f/+
CBPf/+ P300f/f

**Figure S5 related to Figure 2. Cell autonomous loss of neuronal responsiveness after double KAT3 ablation.** Adult *Crebbp*$^{f/f}$*::Ep300*$^{f/f}$ mice with monolateral AAV-Cre-GFP infection in the dentate gyrus were injected intraperitoneally (i.p.) with saline or kainic acid and perfused one hour later during *status epilepticus*. Immunostaining against Fos demonstrates that only the neurons in the non-injected site robustly respond to kainic acid. Scales: 200 µm.

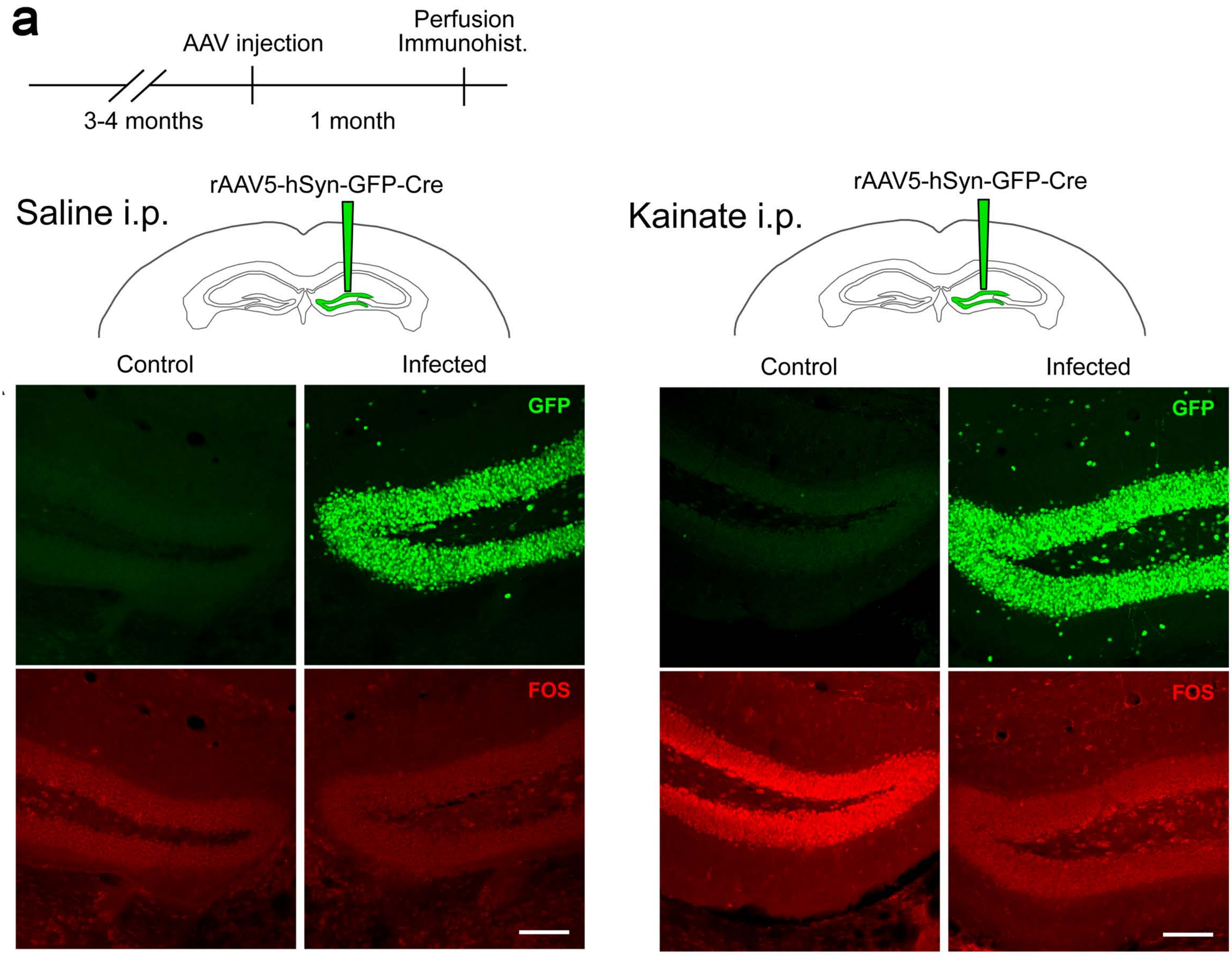
a
Perfusion
Immunohist.
AAV injection
3-4 months
1 month
rAAV5-hSyn-GFP-Cre
Saline i.p.
Control
Infected
GFP
FOS
rAAV5-hSyn-GFP-Cre
Kainate i.p.
Control
Infected
GFP
FOS

**Figure S6 related to Figure 2. KAT3 proteins safeguard the identity of cultured neurons. a.** Immunocytochemistry images showing a robust elimination of CBP and p300 in the GFP-positive neurons of the dKAT3-cKO cultures. Scale bar: 50 µm. **b.** Maintenance of neuronal viability in dKAT3$^{f/f}$::stop$^{f/f}$-tdTomato (tdT) hippocampal neurons infected with LV-CRE up to 27 days post infection. Scale bar: 100 µm. **c.** Staining against neuronal-specific genes NeuroD2 and hippocalcin demonstrate the loss of neuronal identity in dKAT3$^{f/f}$ hippocampal neurons infected with LV-Cre. Scale bar: 50 µm. **d.** Immunostaining against H3K27ac and H2Bac in PNC from dKAT3$^{f/f}$ hippocampi infected with LV-Cre or the control LV-GFP. Scale bar: 50 µm.

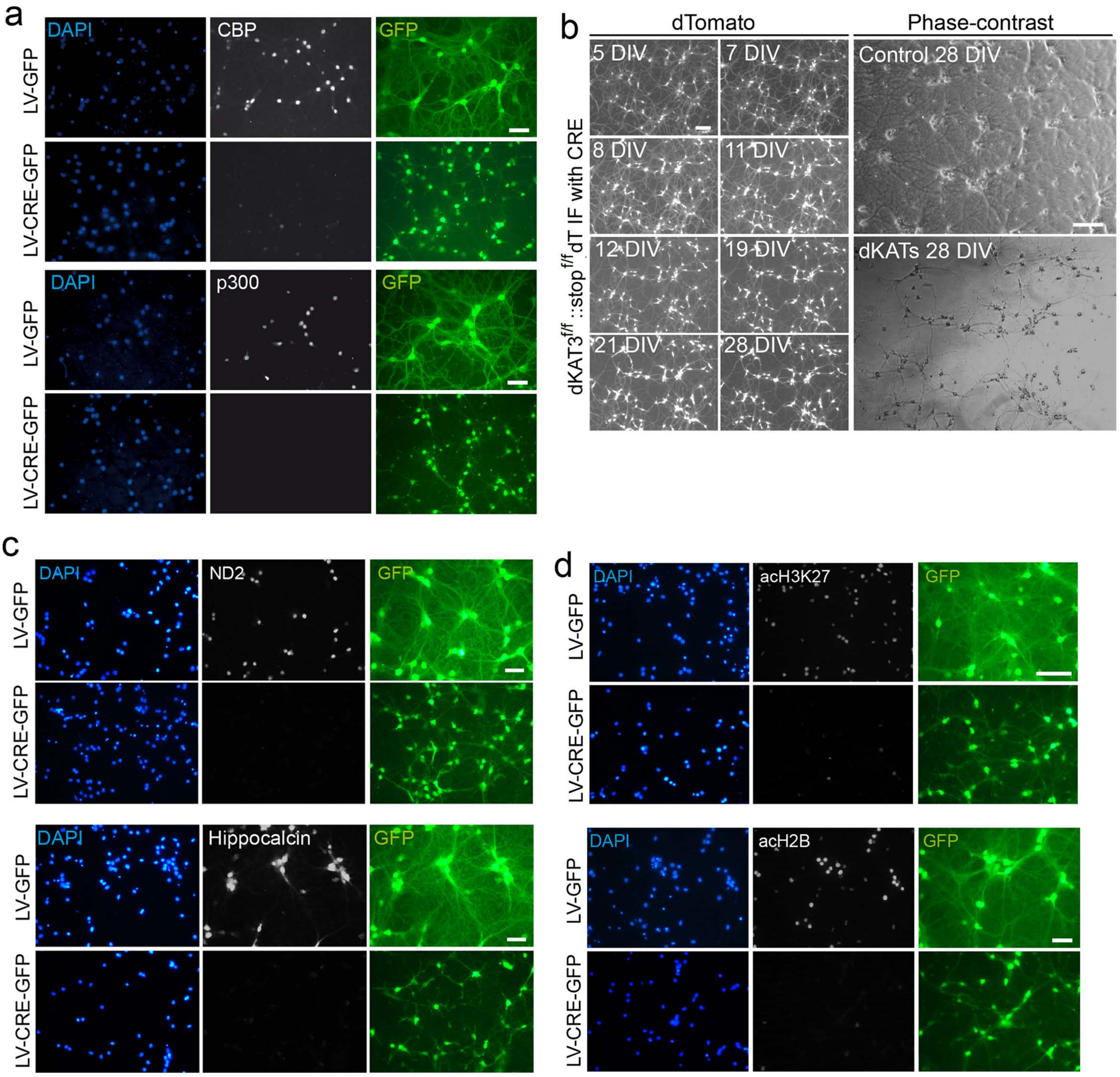
a
DAPI
CBP
GFP
LV-GFP
LV-CRE-GFP
DAPI
p300
GFP
LV-GFP
LV-CRE-GFP
b
dTomato
Phase-contrast
5 DIV
7 DIV
8 DIV
11 DIV
12 DIV
19 DIV
21 DIV
28 DIV
Control 28 DIV
dKATs 28 DIV
dKAT3f/f::stopf/fdT IF with CRE
c
DAPI
ND2
GFP
LV-GFP
LV-CRE-GFP
DAPI
Hippocalcin
GFP
LV-GFP
LV-CRE-GFP
d
DAPI
acH3K27
GFP
LV-GFP
LV-CRE-GFP
DAPI
acH2B
GFP
LV-GFP
LV-CRE-GFP

**Figure S7 related to Figure 3. Loss of neuronal identity examined by single-nucleus RNA-seg. a.** Heat boxes showing the change in expression of genes involved in neurogenesis [40] and the maintenance of neuronal identity [41] in our mRNA-seq data. NSC: Neuronal stem cells; NPC: Neuroprogenitor cells; NB: Neuroblast; CA1 Pyr: CA1 Pyramidal Neurons. **b.** mRNA-seq track for representative genes enriched in NPCs. Scale: 5 kb for *Pax6* and 1 kb for *Fabp7*. **c.** Left: Flow cytometry graphs showing the gate used to isolate singlet nuclei. Right: Sorting accuracy was confirmed by DAPI staining and re-sorting the nuclei. Singlet nuclei positive for DAPI are labelled in red. **d.** UMAP plot showing levels of *Camk2a* transcripts in the control dataset. Note that *Camk2a* expression is restricted to major excitatory neuronal populations such as CA1 and CA3 pyramidal and dentate gyrus granule cells (contours). **e.** Expression of selected subpopulation-specific gene markers in the single-nucleus RNA-seq dataset of control mice. **f-g.** UMAP plots of integrated datasets from the three time points showing the expression level of selected subpopulation-specific genes. The position of the cells expressing gene markers for granule cells (*Glis3*) and CA1 (*Gm2164, Tshz2*) and CA3 (*Trps1*) neurons that are still detected in dKAT3-ifKOs, shifts towards the novel central cluster (**f**). In contrast, the position of cells expressing gene markers for spared populations, such as *Kcnmb2* (interneurons) and *Prr5l* (oligodendrocytes), remain stable (**g**), **h-i.** UMAP plots of integrated datasets showing the disappearance of cells expressing excitatory neuron-specific genes (e.g., *Grin2a*; **h**) and the appearance of a new cluster characterized by the upregulation of non-cell-specific genes (e.g., *Malat1*; **i**).

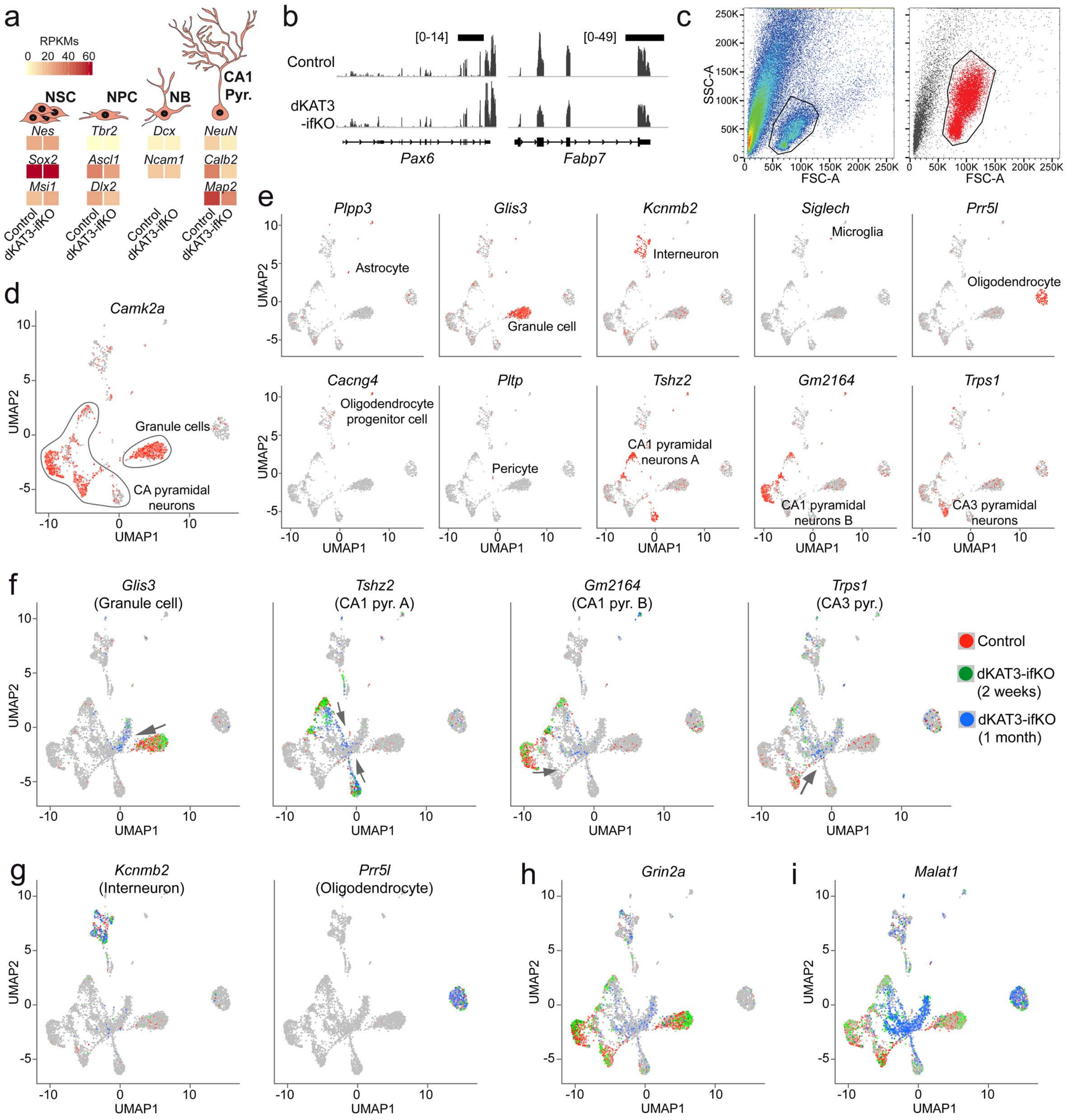
a
RPKMs
0 20 40 60
NSC
NPC
NB
CA1 Pyr.
Nes
Sox2
Msi1
Tbr2
Ascl1
Dlx2
Dcx
Ncam1
NeuN
Calb2
Map2
Control
dKAT3-ifKO
b
[0-14]
[0-49]
Control
dKAT3 -ifKO
Pax6
Fabp7
c
SSC-A
FSC-A
d
Camk2a
Granule cells
CA pyramidal neurons
UMAP1
UMAP2
e
Plpp3
Astrocyte
Glis3
Granule cell
Kcnmb2
Interneuron
Siglech
Microglia
Prr5l
Oligodendrocyte
Cacng4
Oligodendrocyte progenitor cell
Pltp
Pericyte
Tshz2
CA1 pyramidal neurons A
Gm2164
CA1 pyramidal neurons B
Trps1
CA3 pyramidal neurons
f
Glis3 (Granule cell)
Tshz2 (CA1 pyr. A)
Gm2164 (CA1 pyr. B)
Trps1 (CA3 pyr.)
Control
dKAT3-ifKO (2 weeks)
dKAT3-ifKO (1 month)
g
Kcnmb2 (Interneuron)
Prr5l (Oligodendrocyte)
h
Grin2a
i
Malat1

**Figure S8 related to Figure 4. CBP and p300 bind to the same places throughout the mouse genome. a.** ChIP assays demonstrating the specificity of the CBP and p300 antibodies. The *Fos* and *Bdnf* promoters (Pr) are occupied by the KAT3 proteins, while the chosen intergenic location (negative control) is not. **b.** Principal component analysis (PCA) of CBP and p300 ChIP-seqs from control and dKAT3-ifKO. CBP and p300 ChIP-seq samples cluster perfectly depending on the genetic background, indicating they are nearly identical. Genotype explains 78% of the variance. **c.** Left: Overlap of KAT3 peaks CBP and p300 depends on the threshold for Independent Discovery Rate (IDR) used. Here, we maintained a constant IDR of 0.05 for CBP ChIP-seq and calculated the overlap with the p300 ChIP-seq peaks obtained by increasing the IDR in steps of 0.05. Numbers in Venn plots indicate peaks in thousands, lines in the graph indicate % of overlap. At IDR 0.05, most of the p300 peaks (~90%) colocalizes with CBP. With and IDR for p300 of 0.25, most CBP peaks (~90%) co-localizes with p300. Right: The upper profiles show two examples of CBP peaks that are detected or not in p300 profiles, depending on the IDR threshold used. The enrichment for p300 signal over background in clearly observed in both regions.

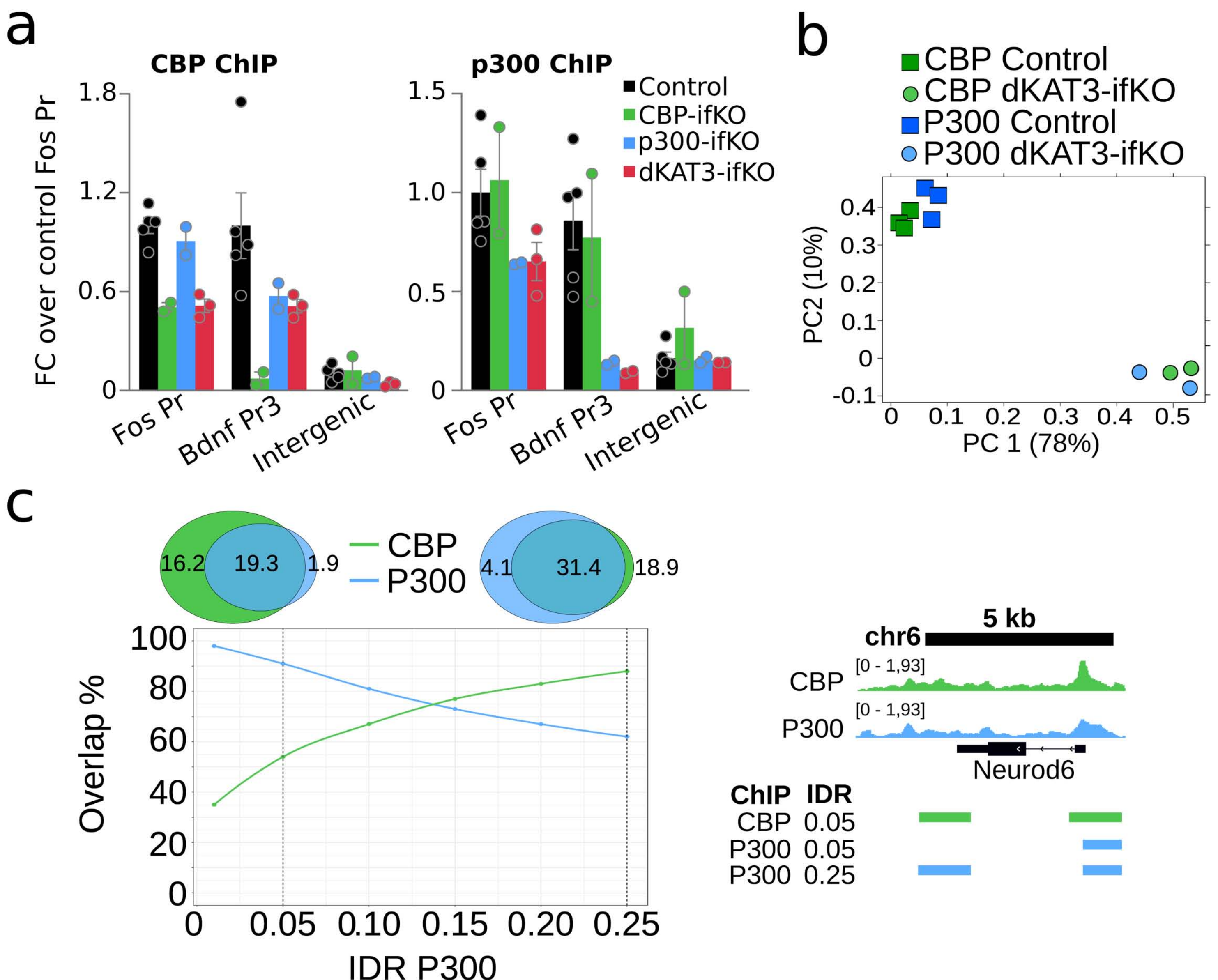
a
CBP ChIP
p300 ChIP
FC over control Fos Pr
1.8
1.2
0.6
0
1.5
1.0
0.5
0
Control
CBP-ifKO
p300-ifKO
dKAT3-ifKO
Fos Pr
Bdnf Pr3
Intergenic
b
CBP Control
CBP dKAT3-ifKO
P300 Control
P300 dKAT3-ifKO
PC2 (10%)
PC 1 (78%)
c
16.2
19.3
1.9
CBP
P300
4.1
31.4
18.9
Overlap %
IDR P300
100
80
60
40
20
0
0.05
0.10
0.15
0.20
0.25
5 kb
chr6
[0 - 1,93]
CBP
P300
Neurod6
ChIP IDR
CBP 0.05
P300 0.05
P300 0.25

**Figure S9 related to Figure 4. Classification of KAT3 peaks according to cell type and gene features. a.** Overlap between ATAC and KAT3 (CBP + P300) peaks. Numbers of regions detected in each screen in thousands. **b.** Metaplots showing the signal for proteins and histone post-translational modifications associated with promoters (H3K4me3, H3K9-14ac and RNAPII) and enhancers (H3K27ac and H3K4me1, as well as CBP and p300 binding), and chromatin accessibility profiles. Data is provided for the sets of neuronal (green), non-neuronal (red) and pancellular (orange) KAT3 peaks. **c.** Distribution of DARs in neuronal and pancellular KAT3 peaks in dKAT3-ifKOs. **d.** Changes in ATAC-seq signal at neuronal and pancellular peaks. **e.** Relationship between loss of accessibility and gene expression for all peaks in dKAT3-ifKO genetic background divided by genomic feature. Both types of change occur almost exclusively in intergenic and intronic regions.

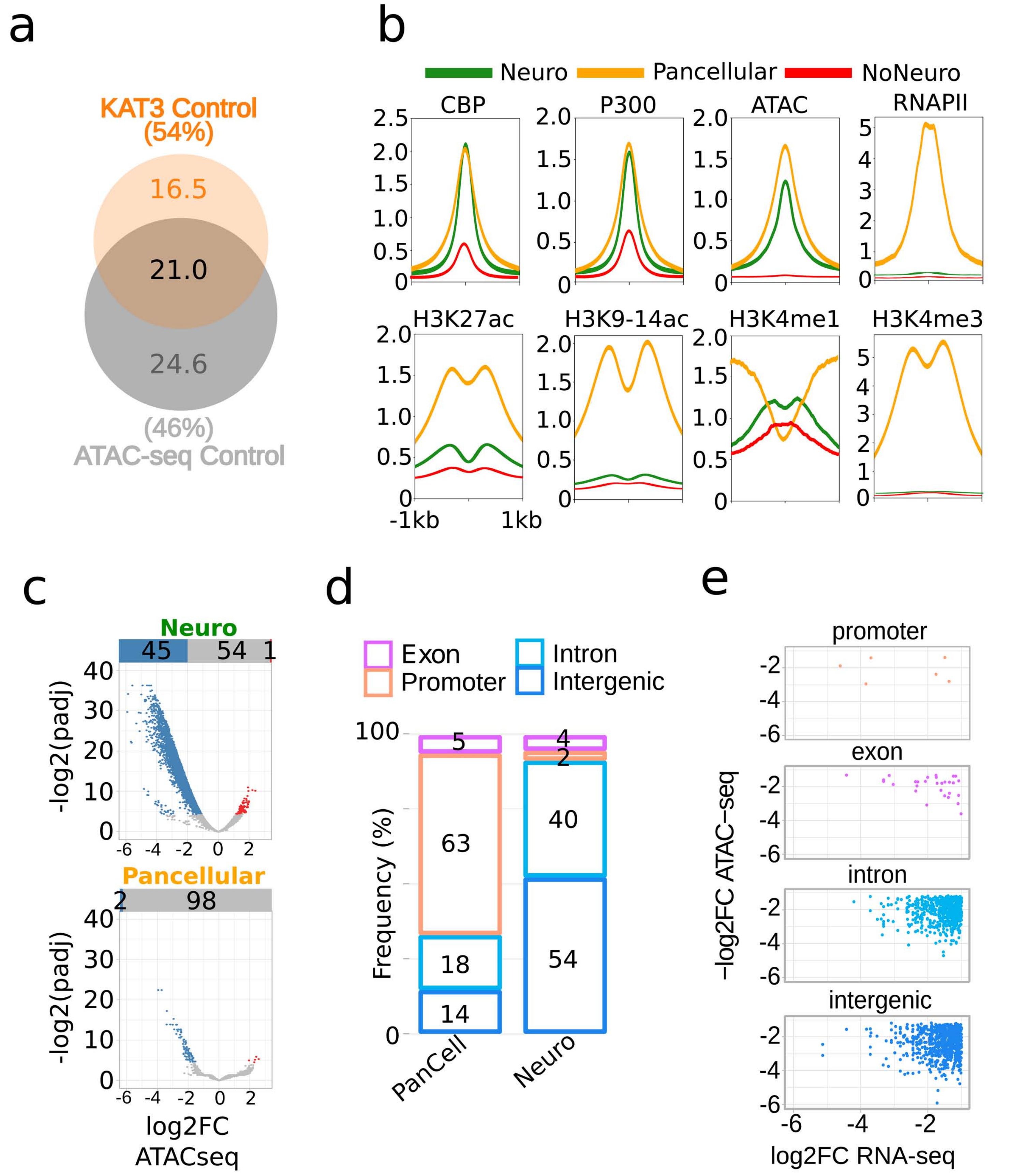
a
KAT3 Control
(54%)
16.5
21.0
24.6
(46%)
ATAC-seq Control
b
Neuro
Pancellular
NoNeuro
CBP
P300
ATAC
RNAPII
H3K27ac
H3K9-14ac
H3K4me1
H3K4me3
-1kb
1kb
c
Neuro
45
54
1
-log2(padj)
Pancellular
2
98
log2FC
ATACseq
d
Exon
Promoter
Intron
Intergenic
Frequency (%)
5
63
18
14
4
2
40
54
PanCell
Neuro
e
promoter
exon
intron
intergenic
-log2FC ATAC-seq
log2FC RNA-seq

**Figure S10 related to Figure 5. H3K27ac hypoacetylation correlates with gene downregulation and overlaps with chromatin interactions specific of differentiated neurons**. **a.** Representative images of immunostaining against different acetylated lysine residues in the chromatin of CA1 pyramidal neurons in brain slices from mice with the indicated genotypes. Scale: 10 μm. **b-c**. Immunostaining for H3K27me3 (b) and H3K9me3 (c) of brain slices of adult *Crebbp$^{f/f}$::Ep300$^{f/f}$* mice 2 months after monolateral AAV-Cre-GFP infection in the dentate gyrus. Scale: 10 μm. **d.** Top panel: Percentage of overlap between H3K27ac and KAT3 peaks in the chromatin of control mice. Percentages are shown for all peaks (Global) and for the cell-specific subsets: neuronal (Neuro), non-neuronal (NoNeuro) and pancellular (Pancell). Middle panel: Reads per kb and million reads (RPKMs) values of the top 20% expressed genes that contain a peak with both H3K27ac and KAT3 signal in the same subsets indicated above. Bottom panel: Gene expression changes in dKAT3-ifKO hippocampus split in the same subsets as above. **e**. Genomic distribution of H3K27ac peaks in wild type mice categorized by genomic feature and distribution of H3K27ac peaks that overlap with neuronal, pancellular and non-neuronal KAT3 peaks. The two rightmost bars show the percentage of H3K27ac regions that loses acetylation categorized by genomic feature, for all peaks and for those that overlap with KAT3 neuronal peaks. **f.** Changes of H3K9,14ac observed in dKAT3-ifKOs hippocampal chromatin. **g.** Eigen values in regions with neuronal enhancer loss in dKAT3-ifKO for the three differentiation stages described in [32]: Embryonic stem cells (ES), neuroprogenitors (NPC) and cortical neurons differentiated *in vitro* (CN). CN vs NPC: p = 0.002; CN vs ES: p = 1.5E-08 (one side Mann-Whitney test). **h.**

Hi-C Eigen-value profiles in ES, NPC, CN and hippocampal neurons from adult mice [24, 32]. Blue: open chromatin. Magenta: closed chromatin. Lower tracks show the positions of the lost neuronal ATAC-seq signal (green), lost neuronal enhancers (cyan) and gene upregulation (red) or downregulation (light blue) in dKAT3-ifKOs.

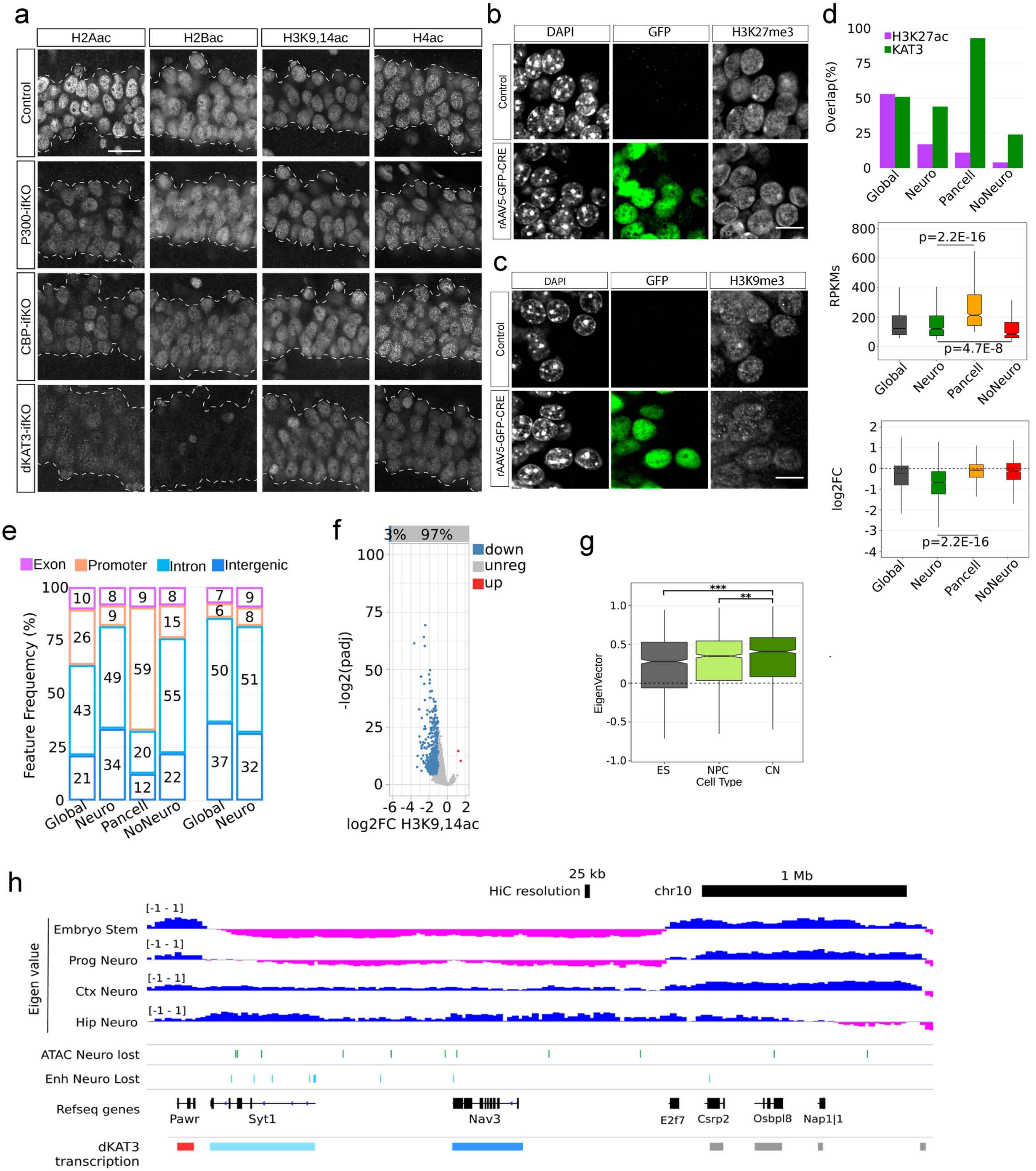

a
H2Aac
H2Bac
H3K9,14ac
H4ac
Control
P300-ifKO
CBP-ifKO
dKAT3-ifKO
b
DAPI
GFP
H3K27me3
Control
rAAV5-GFP-CRE
c
DAPI
GFP
H3K9me3
Control
rAAV5-GFP-CRE
d
H3K27ac
KAT3
Overlap(%)
Global
Neuro
Pancell
NoNeuro
RPKMs
p=2.2E-16
p=4.7E-8
log2FC
p=2.2E-16
e
Exon
Promoter
Intron
Intergenic
Feature Frequency (%)
Global
Neuro
Pancell
NoNeuro
Global
Neuro
f
3%
97%
down
unreg
up
-log2(padj)
log2FC H3K9,14ac
g
EigenVector
ES
NPC
CN
Cell Type
h
25 kb
HiC resolution
1 Mb
chr10
Eigen value
[-1 - 1]
Embryo Stem
Prog Neuro
Ctx Neuro
Hip Neuro
ATAC Neuro lost
Enh Neuro Lost
Refseq genes
Pawr
Syt1
Nav3
E2f7
Csrp2
Osbpl8
Nap1l1
dKAT3 transcription

**Figure S11 related to Figure 5. CBP and p300 are recruited by bHLH transcription factors and regulate their expression. a.** Motif found by a motif enrichment analysis algorithm in the regions with a decreased ATAC-seq signal in dKAT3-ifKO. These regions show a signature indistinguishable from the canonical bHLH TFs binding motif. **b.** Plot showing the expression level and change in expression of bHLHs in dKAT3-ifKO hippocampus versus control mice. Every dot is a single bHLH TF expressed in the hippocampus. Blue – downregulated, red – upregulated, grey – unchanged gene expression. **c.** Left: Footprint of bHLHe22, another bHLH TF expressed in mature excitatory neurons. Right: Footprint of ZNF740, an example of TF detected in both neuronal and pancellular peaks. Values on the y-axis correspond to normalized Tn5 insertions. Values on the x-axis describe the position from the peak in bp. **d.** Overlap between Neurod2 ChIP-seq peaks and bHLH footprints found in ATAC-seq neuro-peaks. Numbers are expressed in thousands. **e.** Representative snapshots of RNA-seq, ATAC-seq, and CBP and H3K27ac ChIP profiles at the gene *Hrk* (left), which encodes a positive regulator of neuronal death strongly downregulated in dKAT3-ifKOs and belong to the category of neuronal-specific genes defined in **Figure 4**. For comparison, we also present the profiles of *Kat5*, which encodes a ubiquitous KAT that is not downregulated in dKAT3-ifKOs and was classified as pancellular gene.

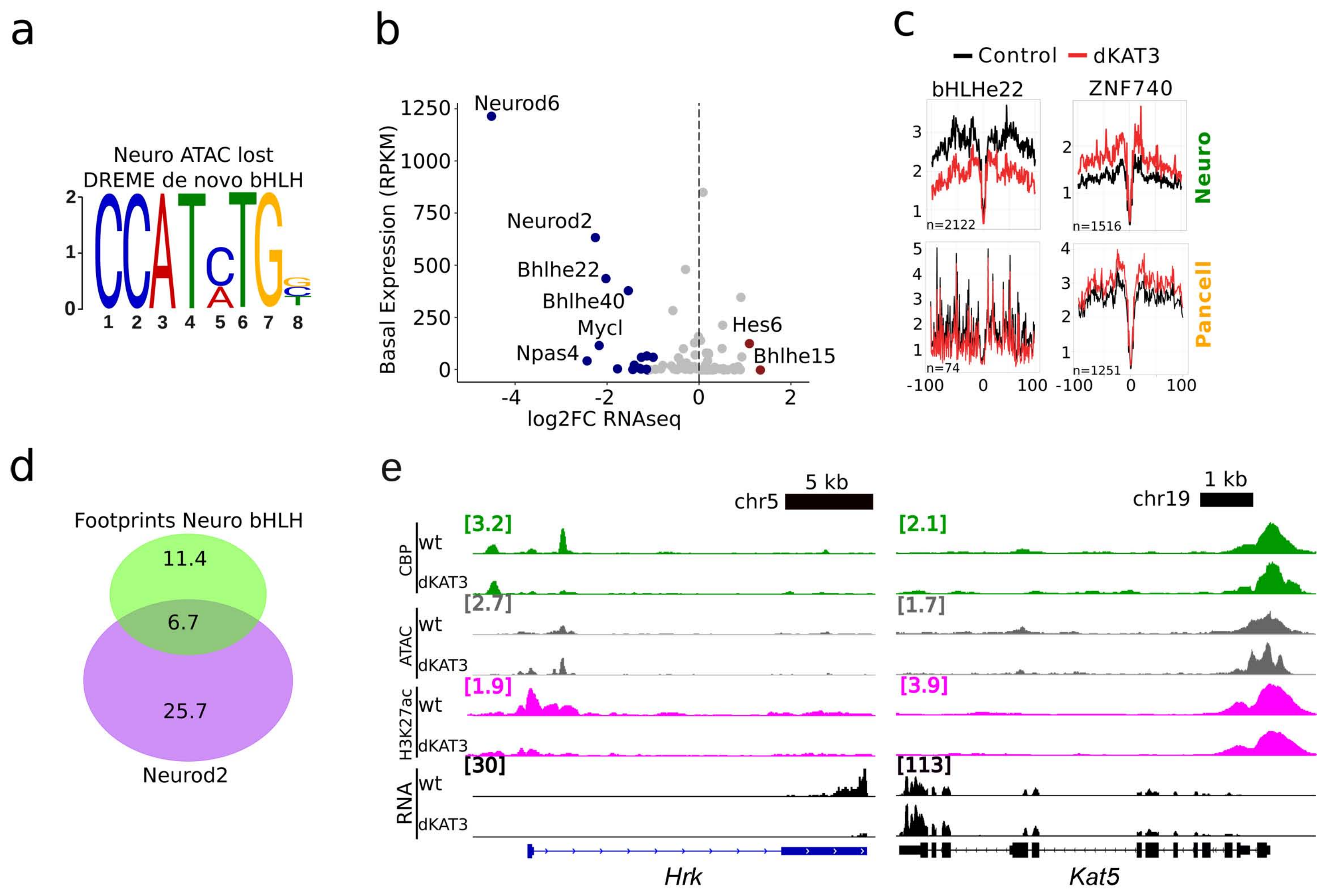

a
Neuro ATAC lost
DREME de novo bHLH
b
Neurod6
Neurod2
Bhlhe22
Bhlhe40
Mycl
Npas4
Hes6
Bhlhe15
Basal Expression (RPKM)
log2FC RNAseq
c
Control
dKAT3
bHLHe22
ZNF740
n=2122
n=1516
n=74
n=1251
Neuro
Pancell
d
Footprints Neuro bHLH
11.4
6.7
25.7
Neurod2
e
5 kb
chr5
1 kb
chr19
CBP
ATAC
H3K27ac
RNA
wt
dKAT3
[3.2]
[2.1]
[2.7]
[1.7]
[1.9]
[3.9]
[30]
[113]
Hrk
Kat5

**Figure S12 related to Figure 8. Overexpression of NeuroD2 does not rescue neuronal-specific transcription deficits in dKAT3-KO neurons.** **a.** Overlap between NeuroD2 peaks [34] and ATAC-seq peaks in the control chromatin. **b.** Gene feature distribution of ATAC-seq enriched regions, NeuroD2 (ND2) ChIP-seq peaks and overlapping regions. **c.** RT-PCR quantifications of neuronal gene markers after LV-driven overexpression of a human version of bHLH transcription factor NeuroD2 (ND2 – *Neurod2*, NeuN – *Rbfox3*) The primers designed for ND2 can distinguish between the human and mouse mRNAs (hND2 and mND2 respectively).

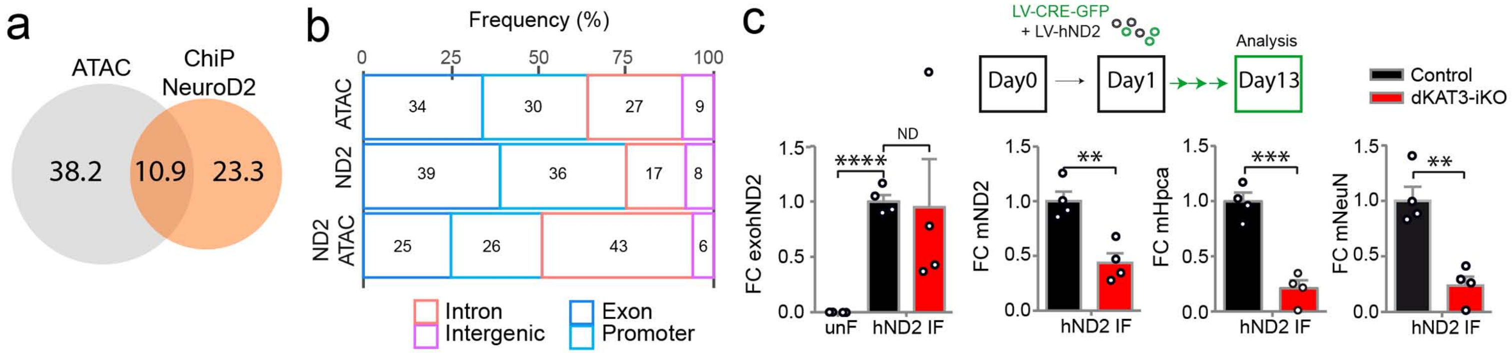
a
ATAC
ChiP NeuroD2
38.2
10.9
23.3
b
Frequency (%)
0
25
50
75
100
ATAC
34
30
27
9
ND2
39
36
17
8
ND2 ATAC
25
26
43
6
Intron
Intergenic
Exon
Promoter
c
LV-CRE-GFP
+ LV-hND2
Analysis
Day0
Day1
Day13
Control
dKAT3-iKO
FC exohND2
****
ND
unF
hND2 IF
FC mND2
**
hND2 IF
FC mHpca
***
hND2 IF
FC mNeuN
**
hND2 IF
1.5
1.0
0.5
0.0

**Table S1 related to Figure 1. Results of SHIRPA screen in single and double KAT3 ifKOs**

| | Test | CBP -ifKO | p300 -ifKO | dKAT3 -ifKO | CreERT2:: Crebbp$^{f/+}$:: Ep300$^{f/f}$ | CreERT2:: Crebbp$^{f/f}$:: Ep300$^{f/+}$ |
|---|---|---|---|---|---|---|
| Spontaneous activity | MW | ns | ns | ns/0.04 | ns/ns | ns/ns |
| Transfer arousal | MW | ns | ns | ns/ns | ns/ns | ns/ns |
| Fear | F | ns | ns | ns/ns | ns/ns | ns/ns |
| Piloerection | F | ns | ns | ns/ns | ns/ns | ns/ns |
| Body position | MW | ns | ns | ns/0.01 | ns/ns | ns/ns |
| Positional passivity | MW | ns | ns | ns/ns | ns/ns | ns/ns |
| Gait | MW | ns | ns | ns/ns | ns/ns | ns/ns |
| Pelvic elevation | MW | ns | ns | ns/ns | ns/ns | ns/ns |
| Tail elevation | MW | ns | ns | ns/ns | ns/ns | ns/ns |
| Visual placing | MW | ns | ns | ns/ns | ns/ns | ns/ns |
| Grip strength | MW | ns | ns | ns/0.03 | ns/ns | ns/ns |
| Wire maneuver | MW | ns | ns | ns/ns | ns/ns | ns/ns |
| Touch escape | MW | ns | ns | ns/ns | ns/ns | ns/ns |
| Trunk curl | F | ns | ns | ns/ns | ns/ns | ns/ns |
| Limb grasping | F | ns | ns | ns/ns | ns/ns | ns/ns |
| Provoked biting | F | ns | ns | ns/ns | ns/ns | ns/ns |
| Irritability | F | ns | ns | ns/ns | ns/ns | ns/ns |
| Aggression | F | ns | ns | ns/ns | ns/ns | ns/ns |
| Vocalization | F | ns | ns | ns/ns | ns/ns | ns/ns |
| Righting reflex | MW | ns | ns | ns/ns | ns/ns | ns/ns |
| Contact righting reflex | F | ns | ns | ns/ns | ns/ns | ns/ns |
| Negative geotaxis | MW | ns | ns | ns/0.03 | ns/ns | ns/ns |
| Pinna reflex | MW | ns | ns | ns/0.02 | ns/ns | ns/ns |
| Corneal reflex | MW | ns | ns | ns/ns | ns/ns | ns/ns |
| Palpebral closure | MW | ns | ns | ns/ns | ns/ns | ns/ns |
| Toe pinch | MW | ns | ns | ns/ns | ns/ns | ns/ns |
| Heart rate | MW | ns | ns | ns/ns | ns/ns | ns/ns |
| Respiration rate | MW | ns | ns | ns/ns | ns/ns | ns/ns |
| Tremor | MW | ns | ns | ns/ns | ns/ns | ns/ns |
| Barbering | MW | ns | ns | ns/ns | ns/ns | ns/ns |
| Body tone | MW | ns | ns | ns/ns | ns/ns | ns/ns |
| Abdominal tone | MW | ns | ns | ns/ns | ns/ns | ns/ns |
| Limb tone | MW | ns | ns | ns/ns | ns/ns | ns/ns |
| Skin color | MW | ns | ns | ns/ns | ns/ns | ns/ns |
| Head morphology | F | ns | ns | ns/ns | ns/ns | ns/ns |
| Lacrimation | MW | ns | ns | ns/ns | ns/ns | ns/ns |
| Salivation | MW | ns | ns | ns/ns | ns/ns | ns/ns |

Table shows results of an adapted SHIRPA test. CBP-ifKO, p300-ifKO, CreERT2:: Crebbp$^{f/+}$::Ep300$^{f/f}$ and CreERT2::Crebbp$^{f/f}$::Ep300$^{f/+}$ do not show any neurological phenotype. dKAT3-ifKO were affected only after tamoxifen (TMX) treatment. The result before and after TMX are separated by a "/" sign. Numbers represent p-value of comparison. Statistical test used for the comparison is indicated in the column "Test": F - Fisher Exact test, MW - Mann-Whitney U test. ns = non-significant.
